# Human STING oligomer function is governed by palmitoylation of an evolutionarily conserved cysteine

**DOI:** 10.1101/2023.08.11.553045

**Authors:** Rebecca Chan, Xujun Cao, Sabrina L Ergun, Evert Njomen, Stephen R. Lynch, Christopher Ritchie, Benjamin Cravatt, Lingyin Li

## Abstract

The anti-viral and anti-cancer STING innate immune pathway can exacerbate autoimmune and neurodegenerative diseases when aberrantly activated, emphasizing a key unmet need for STING pathway antagonists. However, no such inhibitors have advanced to the clinic because it remains unclear which mechanistic step(s) of human STING activation are crucial for potent and context-independent inhibition of downstream signaling. Here, we report that C91 palmitoylation, the mechanistic target of a potent tool compound, is not universally necessary for human STING signaling, making it a poor target for drug development. Instead, we discover that evolutionarily conserved C64 is basally palmitoylated and is crucial for preventing unproductive STING oligomerization in the absence of cGAMP stimulation. The effects of palmitoylation at C64 and C91 converge on the control of intra-dimer disulfide bond formation at C148. Importantly, we show for the first time that signaling-competent STING oligomers are composed of a mixture of two species: disulfide-linked STING dimers that stabilize the oligomer, and reduced STING dimers that are phosphorylated to actuate interferon signaling. Given this complex landscape and cell type specificity of palmitoylation modifications, we conclude that robust STING inhibitors must directly inhibit the oligomerization process. Taking inspiration from STING’s natural autoinhibitory mechanism, we identified an eight amino acid peptide that binds a defined pocket at the inter-dimer oligomerization interface as a proof-of-concept human STING inhibitor, setting the stage for future therapeutic development.

**Summary:** We report that functional STING oligomers require palmitoylation at cysteine 64 and some proportion of reduced dimers, and define the site of autoinhibition that can be targeted to disrupt STING oligomerization and activity.

## Introduction

The innate immune system must be tightly regulated to avoid autoimmunity while responding appropriately to real threats. The STING pathway is a quintessential example where this delicate balance is maintained by many layers of regulatory mechanisms.(*1*) The STING pathway is activated when double-stranded DNA (dsDNA) or DNA:RNA hybrids are detected in the cytosol by cyclic-GMP-AMP synthase (cGAS), which produces the second messenger 2’,3’-cyclic-GMP-AMP (cGAMP) in response.(*2–5*) cGAMP directly binds to STING, a transmembrane receptor that resides on the cytosolic face of the endoplasmic reticulum (ER).(*6*) Activated STING serves as a scaffold to bring the kinase TBK1 to the transcription factor IRF3, leading to phosphorylation, dimerization, and nuclear translocation of IRF3, which then activates the expression and secretion of type I interferons.(*7*, *8*) Activation of the STING pathway has broad antiviral and anticancer effects but also exacerbates many autoimmune and neurodegenerative diseases, necessitating deep understanding of the biochemical basis of STING activation.

Inactive STING exists as a stable dimer auto-inhibited by its C-terminal tail (CTT, aa 340– 379).(*9*) Upon binding to cGAMP, STING undergoes a conformational change that closes the ligand binding domain (LBD) around cGAMP(*10*) and rotates the cytosolic domain 180° relative to its transmembrane domain.(*11*) Release of the CTT from its autoinhibitory binding site exposes STING’s oligomerization interface, allowing it to form disulfide-stabilized oligomers via C148.(*9*) The formation of STING oligomers then initiates an irreversible sequence that includes translocation from the ER to the Golgi and palmitoylation at the transmembrane cysteines C88 and C91, while the released CTT is arrayed on the STING oligomer to scaffold multivalent interactions between TBK1 and IRF3 for downstream signaling.(*7*, *8*) However, no investigational new drugs have advanced into clinical trials 15 years after discovery of STING due to a lack of enabling biochemical and molecular mechanisms. For example, CTT and palmitoylation are not visible and disulfide bonds are not present due to the reducing conditions used in existing STING structures.(*7*, *12–15*) In addition, the efficacy and mode of action of recently reported inhibitor tool compounds, such as C91-targeting palmitoylation inhibitor H-151(*16*) and C148-targeting BB-Cl-amidine (*17*), remain to be elucidated.

## Results

### STING C148-mediated disulfides form rapidly upon cGAMP induction to stabilize STING oligomers

We and others have previously reported the importance of cysteine 148-mediated disulfide bond (C148) in STING signaling, (*9, 17–19*) leading to the hypothesis that a disulfide linkage between adjacent STING dimers stabilizes STING oligomers. Cryo-EM structures of cGAMP-bound STING have since shown that the C148 residues between adjacent STING dimers are too distant, while the distance between C148 residues within the same STING dimer (intradimer) is within the range of a disulfide linkage,(*12*, *13*), (*20*) raising the question of how such intradimer disulfides might influence STING oligomer formation, stability, and signaling.

We first sought to benchmark the timing of intradimer C148 disulfide formation against other known STING signaling events such as palmitoylation, oligomerization, ER-to-Golgi translocation, and the phosphorylation of STING, TBK1 and IRF3. Using a blue-native PAGE (BN-PAGE) assay in either 293T cells expressing exogenous STING or Hela Kyoto cells expressing an inducible STING construct, we found that oligomerization occurs rapidly, becoming saturated within 10 minutes of cGAMP addition and persisting across the entire time course of 30 min or 2 hours, respectively (**Figure 1A, S1A**). The C148A mutant STING formed similar levels of oligomers at 5 min and 10 min, but the oligomeric band intensity weakened at 30 min. These data also suggest that C148 disulfide bonds, while not necessary for oligomerization, increase oligomer stability. STING disulfide bond formation saturated at 5 min after cGAMP addition, suggesting that disulfide bond formation precedes oligomerization (**Figure 1B, S1B**).

**Figure 1.**
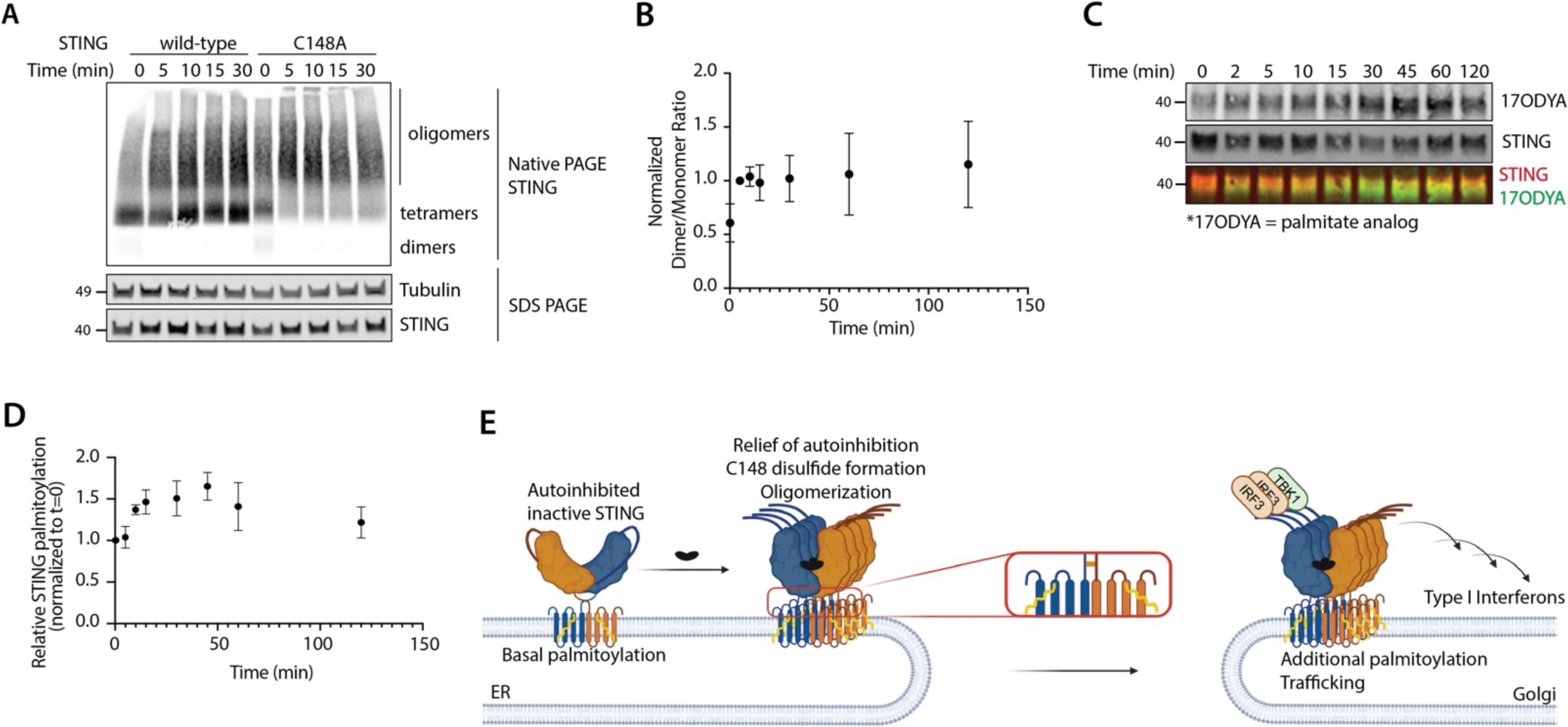
STING is basally palmitoylated and cysteine 148 disulfides occur rapidly to stabilize oligomers. **(A)** Timecourse of STING oligomerization by Blue-Native PAGE of HEK293T cells transfected with STING variants. Cells were transfected overnight with the indicated variant and treated with 100 µM cGAMP for the indicated durations. **(B)** Quantification of dimer to monomer band ratio overtime from non-reducing SDS-PAGE. Samples are HEK293T cells transfected with WT-STING and treated with 100 µM cGAMP for the indicated times. All ratios normalized to *t*=5 min and averaged over three individual experiments. **(C)** Timecourse of STING palmitoylation in Hela Kyoto cGAS KO cells stably expressing STING-FLAG using clickable palmitate analog, 17-ODYA. Cells were pre-treated with 17-ODYA for 1.5 hours, then with 100 µM cGAMP for the indicated durations before palmitoylation was assessed. **(D)** Quantification of palmitoylation assay from (C). All ratios normalized to *t*=0 min and averaged over four individual experiments. **(E)** Model of our current understanding of the STING signaling cascade upon cGAMP addition.

In agreement with previous literature,(*21*, *22*) we confirmed by fluorescence confocal microscopy that STING is primarily ER-localized in the absence of cGAMP and only translocates to the Golgi after addition of cGAMP (**Figure S1C**). The majority of STING translocates to the Golgi approximately 30 minutes after cGAMP addition (**Figure S1D**), and isolation of COP-II vesicles—marking anterograde ER-to-Golgi transport—confirmed that STING is entirely oligomerized during ER-to-Golgi trafficking (**Figure S1E, F**). Together, these data confirm that oligomerization of human STING takes place at the ER prior to Golgi translocation. Since C148A mutant oligomers start to disintegrate before translocation is complete (**Figure 1A**) and this mutant is known to have impaired trafficking ability,(*9*) our results suggest that this intradimer disulfide bond is important for stabilizing STING oligomers sufficiently to survive trafficking.

Finally, we monitored STING palmitoylation over the same time course using the alkynylated palmitate substrate, 17-ODYA, followed by visualization by click addition of an azide-labeled fluorescent dye (**Figure 1C**).(*23*) Although STING is known to be palmitoylated after cGAMP stimulation, basal palmitoylation has never been reported. Furthermore, DHHC enzymes that palmitoylate STING were reported to be Golgi-localized, suggesting that palmitoylation occurs after trafficking.(*24*) We were therefore surprised to observe basal palmitoylation before cGAMP stimulation (at *t* = 0), which increased over time concurrently with translocation to twice the basal level (**Figure 1D**). Taken together, STING disulfide bond formation and oligomerization is the first step after cGAMP induction and occurs before trafficking to the Golgi. Palmitoylation occurs both basally at the ER and in a cGAMP-inducible fashion concomitant with Golgi translocation (**Figure 1E**).

### Blocking palmitoylation at C91 inhibits human STING disulfide bond formation and oligomerization in a context-dependent manner

Given the surprising finding that human STING palmitoylation occurs basally in the absence of cGAMP stimulation, we sought to investigate the role of both basal and cGAMP-induced palmitoylation in human STING signaling. In mouse STING, C91 and neighboring C88 have been shown to be palmitoylated upon STING activation. These two cysteines lie near the cytosol/membrane interface of STING, which would presumably allow them to be targeted by nearby palmitoylation enzymes (**Figure 2A**) and this post-translational modification was shown to be important for downstream signal transduction. (*24*) A recent study showed that C88/91 palmitoylation facilitates higher-order clustering of STING into lipid-rich regions of the Golgi.(*24*, *25*) A valuable tool compound to block STING palmitoylation is H-151, a cysteine-reactive compound that targets C88 and C91 in the transmembrane region of both mouse and human STING.(*16*) H-151 has shown great efficacy at blocking STING activation in the THP-1 human cell line as well as in murine cell lines and mouse models. We tested H-151 in THP1 and U937 cells and observed inhibition of STING signaling (**Figure 2B**). By RNA-seq analysis, H-151 treated U937 cells exhibited altered expression of most of the genes that were also altered in STING^-/-^ cells, further suggesting the potency of this molecule towards STING inhibition in this cell line (**Figure S2A**). H-151 treatment also inhibited C148 disulfide bond formation by more than 50% (**Figure 2C, S2B**) and destabilized oligomers by a similar magnitude to the C148A mutant (**Figure 1A, 2D**), pointing to a previously undiscovered allosteric relationship between C148 and C91.

**Figure 2.**
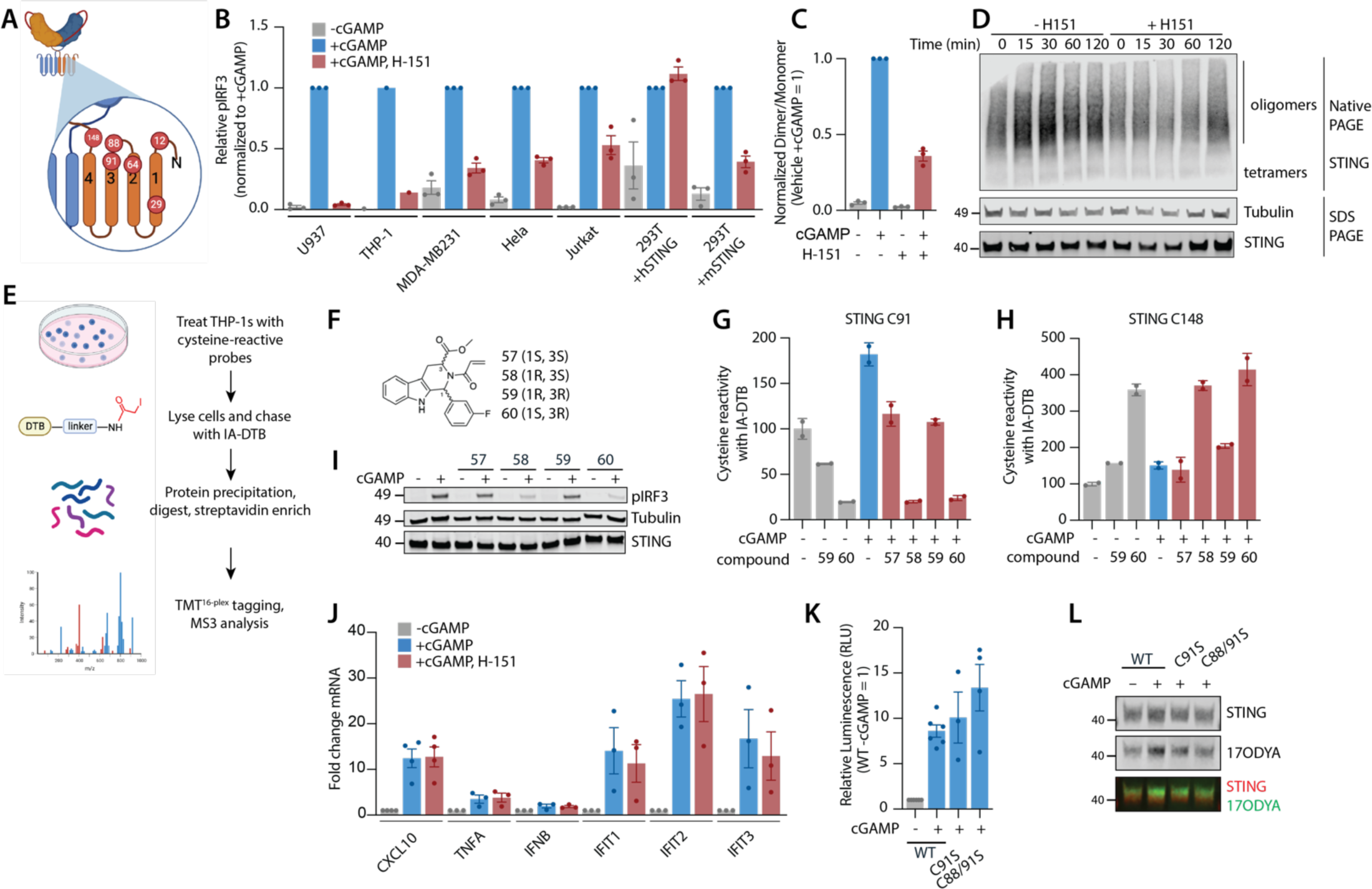
Blocking palmitoylation at C91 allosterically affects C148 and inhibits STING in a context-dependent manner. **(A)** Cartoon depiction of STING’s transmembrane domain and location of all cysteines in the transmembrane and linker regions. **(B)** Relative pIRF3 levels of cells pre-treated with 5 µM H-151, then stimulated with 100 µM cGAMP for an additional 2h. All levels were normalized to their corresponding cGAMP treatment alone, amongst a panel of six cell lines. **(C)** Quantification of dimer to monomer band ratio in U937 cells pre-treated with either DMSO or 5 µM H-151 for 2 h, followed by 100 µM cGAMP stimulation for an additional 2h. All ratios normalized to vehicle +cGAMP condition and averaged over three individual experiments. **(D)** Blue-Native PAGE assay of U937 cells pre-treated with or without H-151 followed by 100 µM cGAMP for the indicated durations. **(E)** Schematic of whole-proteome reactive cysteine profiling workflow. **(F)** Chemical structures of four cysteine-reactive stereoisomers, compounds 57–60. **(G)** Changes in C91 reactivity relative to DMSO control upon indicated compound and cGAMP treatment conditions in THP-1 cells. Cells were treated with 10µM compound alone or both compound and 100µM cGAMP concurrently **(H)** Changes in C148 reactivity relative to DMSO control upon different compound and cGAMP treatment conditions in THP-1 cells. **(I)** U937 cells pre-treated with 10 µM of each compound 57–60 shown in (F) for 2h, followed by stimulation with cGAMP. **(J)** RT-qPCR of various cytokine mRNA levels in peripheral blood mononuclear cells (PBMCs) under indicated conditions. **(K)** IFNβ-luciferase reporter corresponding to interferon response in HEK293T-Dual reporter cells transfected with indicated STING variants followed by cGAMP stimulation for 20 h. **(L)** Palmitoylation assay of HEK293T cells transfected with indicated STING variants followed by cGAMP stimulation for 2 h.

To further characterize the crosstalk between C148 disulfide bond formation and C91 palmitoylation, we employed a broad whole-proteome reactive cysteine profiling platform. Additionally, by using all stereoisomers for each cysteine-reactive probe, we can better distinguish mechanistic details and druggability of each cysteine(*26*) (**Figure 2E**). Probe-reactive cysteines were identified by their subsequent reduced reactivity to a pan-reactive iodoacetamide probe (IA-DTB). We identified two additional compounds **58** and **60** that are enantiomers of each other (**Figure 2F**) that both conjugate onto human STING in THP-1 cells at C91 (**Figure 2G**). In addition, engagement of C91 by either **58** or **60** led to increased reactivity of C148, suggesting that C148 becomes more available when C91 is liganded, consistent with our observation that H-151 reduced C148 disulfides (**Figure 2H**). Also like H-151, both enantiomers inhibited STING signaling in U937 cells with IRF3 phosphorylation (pIRF3) as a readout, suggesting C91-reactive compounds effectively inhibit STING in these cells (**Figure 2I**). In aggregate, C91 palmitoylation inhibitors block STING signaling by allosterically controlling C148 disulfide bond formation. However, the fact that compounds **58** or **60** as stereoisomers both inhibited human STING activity suggests a lack of stereochemical discrimination at the region surrounding C91, which limits the potential of this palmitoylation site as a target of STING inhibitor development (see Discussion). In further support, H-151-treated cells exhibited a transcriptome distinct from untreated cells even in the absence of STING, suggesting non-specific effects (**Figure S2A**).

Despite the potency of H-151 in inhibiting STING signaling in THP-1 and U937 cells, both of which are cancerous monocytic cell lines, we discovered that its efficacy is more varied across other human cell lines containing endogenous STING even at micromolar concentrations (**Figure 2B)**. Similarly, compound **58** and **60** only partially inhibited IRF3 phosphorylation and STING oligomerization (**Figure S2C**). In addition, H-151 did not inhibit STING signaling in primary peripheral blood mononuclear cells (PBMCs) freshly isolated from donors (**Figure 2J**). All three compounds did not block cGAMP-induced C148 disulfide bond formation in HEK293T cells exogenously expressing human STING (**Figure S2D**). Most tellingly, H-151 effectively inhibited HEK293T cells exogenously expressing mouse STING, suggesting its efficacy is in fact species specific (**Figure 2B**). The lack of efficacy of H-151 against human STING is not due to a lack of reactivity, as an alkyne-bearing H-151 analog was able to label STING in a C91-dependent manner (**Figure S2E**). This puzzling inconsistency of target engagement of C91 without inhibition suggest that C91 palmitoylation may not be broadly necessary for human STING signaling outside a few malignant monocyte cell lines. Indeed, C88S, C91S, or C88S/C91S STING mutants were as responsive to cGAMP as WT STING, indicated by pIRF3 induction and interferon production (**Figure S2F, 2K**). Intriguingly, serine mutants of C88 and/or C91 were still metabolically labeled by the palmitate analog 17-ODYA upon cGAMP stimulation (**Figure 2L, S2G**), indicating that palmitoylation may also occur elsewhere in human STING.

### Basal palmitoylation of C64 is required for active STING oligomer conformation and signaling

To systematically define all possible palmitoylation sites in human STING and their effects on STING activation, we first created a STING mutant where all five cysteines in the transmembrane domain (**Figure 2A**)—considered to be plausible palmitoylation sites— were mutated to serine (C12S/C29S/C64S/C88S/C91S, hereafter referred to as 5C5S). While STING 5C5S was still stably expressed in HEK293T cells, it exhibited little to no palmitoylation and was completely inactive, suggesting that palmitoylation is restricted to these five cysteines and is required for STING signaling (**Figure 3A–D**). Restoring only C91 (5C5S+91) resulted in a marginal increase in palmitoylation upon cGAMP stimulation, but did not restore STING activity (**Figure 3A–D**). Interestingly, with restoration of only C64 (5C5S+64), we observed only basal palmitoylation, which did not increase further with cGAMP stimulation. Furthermore, C64 alone rescued STING activity to comparable levels as WT STING following cGAMP stimulation, implying that basal palmitoylation at C64 is necessary for human STING activity. Indeed, the single point mutant STING C64S was unresponsive to cGAMP. Of note, the STING C64S mutant was still basally palmitoylated, suggesting that palmitoylation can also occur basally at other cysteines in the absence of C64. However, only palmitoylation at C64 is necessary for human STING activation.

**Figure 3.**
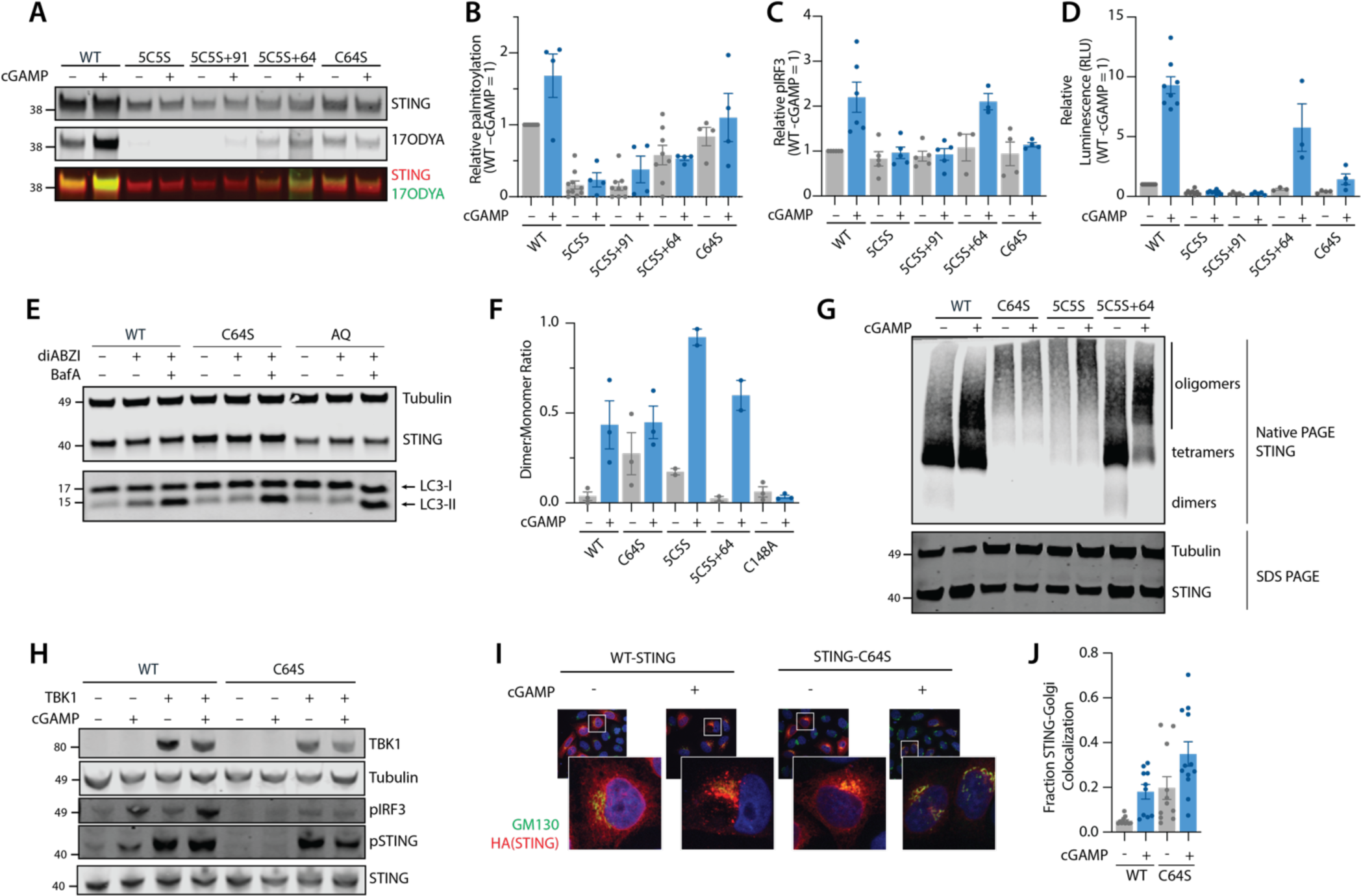
Palmitoylation at C64 alone is crucial for the functional STING oligomer assembly. **(A)** Palmitoylation assay of HEK293T cells transfected with indicated STING variants followed by cGAMP stimulation for 2 h. **(B)** Quantification of palmitoylation assays from (A). **(C)** Relative pIRF3 levels of HEK293Ts transfected with indicated STING variants followed by cGAMP stimulation. **(D)** IFNβ-luciferase reporter corresponding to interferon response in HEK293T-Dual reporter cells transfected with indicated STING variants followed by cGAMP stimulation for 20 h. **(E)** LC3-II levels as a marker for autophagy in HEK293T cells transfected with indicated STING variants, pre-treated with DMSO or 500nM bafilomycin A (BafA) for 2 h before stimulation with 100nM diABZI for an additional 2 h. BafA conditions were included to verify that accumulation of LC3-II was a result of increased autophagic flux rather than lysosomal dysfunction, since inhibition of lysosomal acidification by BafA should result in further increase in LC3-II. **(F)** Quantification of dimer to monomer band ratio in HEK293T cells transfected with indicated STING variants and stimulated with cGAMP. All ratios are averaged over three individual experiments. **(G)** Blue-Native PAGE assay of HEK293T cells transfected with STING variants followed by cGAMP stimulation. **(H)** pIRF3 and pSTING levels of HEK293T cells co-transfected with either WT-STING or STING-C64S mutant along with TBK1-FLAG, followed by cGAMP stimulation. **(I)** Confocal microscopy of Hela Kyoto cGAS KO cells expressing WT-STING or STING-C64S. GM-130 was used as the Golgi marker. **(J)** Quantification of STING–Golgi colocalization from images in (I).

Intriguingly, while C88 and C91 are conserved in many mammals, C64 is much more ancient as it is highly conserved among metazoan STING orthologs down to the sea anemone (**Figure S3A**). In more ancient organisms, STING does not signal via the interferon pathway, but induces autophagy as an immune response.(*27*) The conservation of C64 thus suggests its potentially conserved role in STING-dependent autophagy induction. To test this hypothesis, we investigated autophagy in various STING mutants. Consistent with previous literature,(*28*) WT but not the trafficking-deficient mutant Q273A/A277Q (STING-AQ) was able to induce autophagy upon stimulation in cells, shown by increased LC3 lipidation. C64S was similarly unable to induce autophagy, suggesting that C64 is also important for the autophagy function of STING (**Figure 3E**).

To evaluate why basal C64 palmitoylation is important for human STING signaling, we probed each step of STING signaling starting from disulfide bond formation to determine at what point the C64S mutant loses activity. In contrast to WT which formed disulfide bonds only in the presence of cGAMP, a significant proportion of STING C64S and 5C5S were basally present as the disulfide band prior to cGAMP stimulation (**Figure 3F, S3B**). Strikingly, the STING 5C5S+64 mutant behaved similarly to WT STING and only formed C148 disulfide-linked dimers upon cGAMP stimulation (**Figure 3F, S3B**). These results demonstrated that basal C64 palmitoylation allosterically prevents C148 from forming disulfide bonds prior to cGAMP stimulation.

Since we showed above that C148 disulfide bonds stabilize oligomer formation, we asked whether the C64S mutant, which forms C148 disulfides without stimulation, is constitutively oligomerized. Indeed, BN-PAGE revealed that while WT STING oligomerization only occurs following addition of cGAMP, C64S already formed oligomers in the basal, unstimulated state, and this level did not increase with cGAMP induction (**Figure 3G**). In addition, these C64S oligomers were at noticeably higher molecular weights than active WT oligomers. STING C64S, like WT, can become phosphorylated following TBK1 co-expression and trafficked to the Golgi upon stimulation (**Figure 3H–J**). Co-immunoprecipitation experiments also indicated that this mutant retained the ability to bind TBK1 and IRF3 (**Figure S3C, S3D**). However, IRF3 phosphorylation by TBK1 was impaired even under conditions of TBK1 overexpression, pinpointing this mutant’s loss of activity (**Figure 3H**). We propose that C64S may form an inactive oligomer conformation that is not able to facilitate the productive interaction between TBK1 and IRF3.

To gain additional insight into the nature and driving force of STING C64S oligomers, we evaluated the effects of installing the C64S mutation into a previously identified STING LBD double mutant, A273Q/Q277A (STING-AQ), shown to abolish STING oligomerization and subsequent signaling (Shang et al., 2019). Surprisingly, we found that addition of C64S to the STING-AQ mutant background (C64S-AQ) resulted in oligomerization even in the absence of cGAMP, similar to the C64S mutant alone (**Figure S3E**). We next treated STING-C64S expressing cells with a STING agonist, compound C53, which has previously been shown to bind in the transmembrane domain to promote productive transmembrane interactions between STING dimers.(*13*, *29*) Although compound C53 did not inhibit cGAMP-independent C64S oligomer formation, it did partially rescue STING activity, suggesting that small molecule binding at the transmembrane domain that facilitates appropriate transmembrane interactions could partially compensate for the lack of palmitoylation at C64 (**Figure S3F, S3G**). Together, our results suggest that C64 palmitoylation is crucial for maintaining the activation competent transmembrane domain conformation of human STING and preventing excessive disulfide bond formation.

### Active STING oligomers consist of both disulfide-stabilized and reduced STING dimers

Next, we aimed to reconcile our findings that C148 disulfide bonds are required for STING signaling, while excessive or early C148 disulfide formation (as in the C64S mutant) is detrimental. Strikingly, while all C148-containing STING variants can be phosphorylated upon cGAMP stimulation, we found that the pSTING species was not disulfide-bonded in variants capable of downstream interferon signaling (WT, C91S, 5C5S+64), whereas it was exclusively disulfide-bonded in the inactive mutants (C64S, 5C5S, 5C5S+91) (**Figure 4A, 4B**). Altogether, this non-reducing PAGE analysis allowed us to show for the first time that signaling-competent human pSTING is reduced at C148 in the context of oligomers. Furthermore, since all inactive mutants we tested also lack C64, we conclude that palmitoylation at C64 is required for the formation of this signaling species.

**Figure 4.**
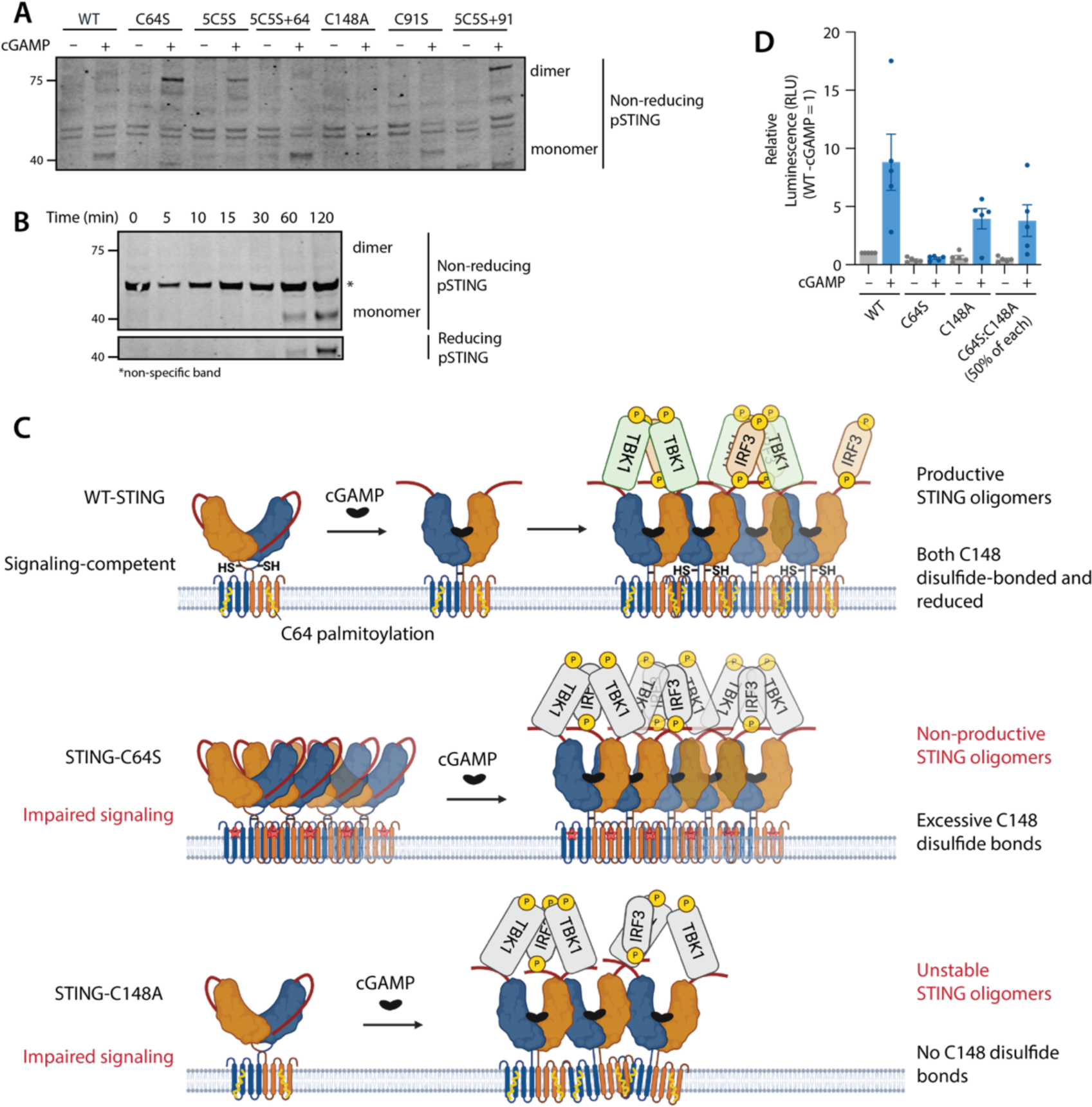
A revised model of STING activation. **(A)** Non-reducing SDS-PAGE of pSTING from HEK293T cells transfected with indicated STING variants followed by 100µM cGAMP stimulation. **(B)** Timecourse of pSTING formation overtime in HEK293T cells transfected with WT-STING and treated with cGAMP for indicated times. **(C)** New model of STING activation: cGAMP binding induces disulfide formation and oligomerization, with the former helping to stabilize oligomers for trafficking. TBK1 docks onto STING and preferentially phosphorylates reduced STING dimers, allowing IRF3 to dock at these sites. Preventing C64 palmitoylation (STING-C64S) results in too much C148 disulfide-linked dimers, which leads to inactive STING oligomers that cannot scaffold appropriate TBK1-IRF3 interactions. When STING cannot form disulfide-linked dimers at all (STING-C148A), this results in unstable STING oligomers that can no longer be trafficked efficiently to the Golgi. **(D)** IFNβ-luciferase reporter corresponding to interferon response in HEK293T-Dual reporter cells transfected with indicated STING variants followed by cGAMP stimulation for 20 h. For the condition where both STING-C64S and STING-C148A are expressed, each variant was expressed in equal amounts.

Together, these findings led us to propose a new model that the signaling-competent STING oligomer is not composed of homogenous STING dimers, but a mixture of disulfide-linked and reduced STING dimers (**Figure 4C**). At the early stages of STING activation, we propose that cGAMP binding induces STING oligomerization, and disulfide formation at C148 stabilizes the oligomer sufficiently for trafficking. STING C148A oligomers, which lack such stabilization, dissociate before or during the trafficking stage. Upon trafficking to the Golgi, TBK1 docks onto disulfide-linked STING dimers and reduced STING dimers equally well, but preferentially phosphorylates the STING CTT on reduced dimers—which we suspect is likely due to a specific conformational requirement—and only this reduced pSTING species allows for IRF3 phosphorylation and downstream signaling. The balance between oxidized and reduced forms of C148 is crucial and is allosterically inhibited by C64 palmitoylation but promoted by C91 palmitoylation in some contexts. The fine balance between the two forms is likely also maintained by the close proximity between intradimer C148 residues and the reducing conditions of the cytosol where C148 is facing.

Our model would predict that artificially mixing oxidized and reduced human STING dimers would be sufficient to rescue STING signaling. To test this, we expressed equal amounts of two mutants that are incompetent for downstream signaling on their own: C64S, which yields excessive disulfide bonds, and C148A, which cannot form disulfides. Strikingly, as predicted by our model, co-expression of these two mutants in equal proportion (while maintaining the same total amount of STING) resulted in an active signaling complex (**Figure 4D**). We therefore conclude that signaling-competent human STING oligomers must be composed of both disulfide-bound and reduced C148 STING dimers.

### Inhibiting STING oligomerization by targeting its autoinhibitory protein-protein interaction interface

The crucial role that STING oligomer composition plays in downstream signaling outputs—fine-tuned by cysteine posttranslational modifications that can vary across species and cell types—points to oligomerization as the key process to inhibit for any robust and context-independent STING inhibitor. We reasoned that blocking STING oligomerization by mimicking its CTT autoinhibitory mechanism(*9*) would broadly inhibit STING signaling in different cellular contexts with a low risk of off-target cellular effects.

In order to explore this inhibitory strategy, we first sought to identify the minimum autoinhibitory motif by splitting the 40-amino acid CTT into three segments (CTTseg1, aa 340–353; CTTseg2, aa 354–366; CTTseg3, aa 367–379, **Figure 5A**) and expressed each HA-tagged segment as bait to co-immunoprecipitate STING. We observed that CTTseg1 and CTTseg3 were able to bind to STING, but CTTseg2 was not (**Figure 5B**). Only CTTseg3 overexpression was able to inhibit cGAMP-induced STING activity in 293T-Dual cells also transiently expressing WT STING (**Figure S4A**). Using confocal microscopy, we also observed that CTTseg3, but not CTTseg1, was able to inhibit the formation of STING puncta (**Figure S4B**), demonstrating that CTTseg3 is the relevant segment in the inhibition of STING oligomerization. We further divided CTTseg3 into two smaller segments and observed that the first half, a seven-residue unit (MEKPLPL, hereon referred to as CTT7, aa 368–374), but not the latter half (PLRTDFS, hereon referred to as nCTT7, aa 373-379) was sufficient to significantly inhibit cGAMP-induced interferon signaling (**Figure 5C**). Likewise, CTT7 but not nCTT7 abolished the formation of STING puncta upon cGAMP treatment, indicating inhibition of STING oligomerization (**Figure 5D**). CTT7 binds the STING cytosolic domain with a moderate affinity of 10.7 µM (**Figure 5E**) and an excess of CTT7 disrupted the cGAMP-induced formation of higher molecular weight species of purified STING cytosolic domain in vitro (**Figure S4C**). Finally, treatment of HEK293T cells with CTT7 peptide disrupted both basal, overexpression-induced STING oligomerization and cGAMP-stimulated oligomerization within 15 min of cGAMP stimulation (**Figure S4D**). Together, we show that CTT7 is a minimal peptide that inhibits STING signaling by directly binding the STING cytosolic domain and preventing STING oligomerization.

**Figure 5.**
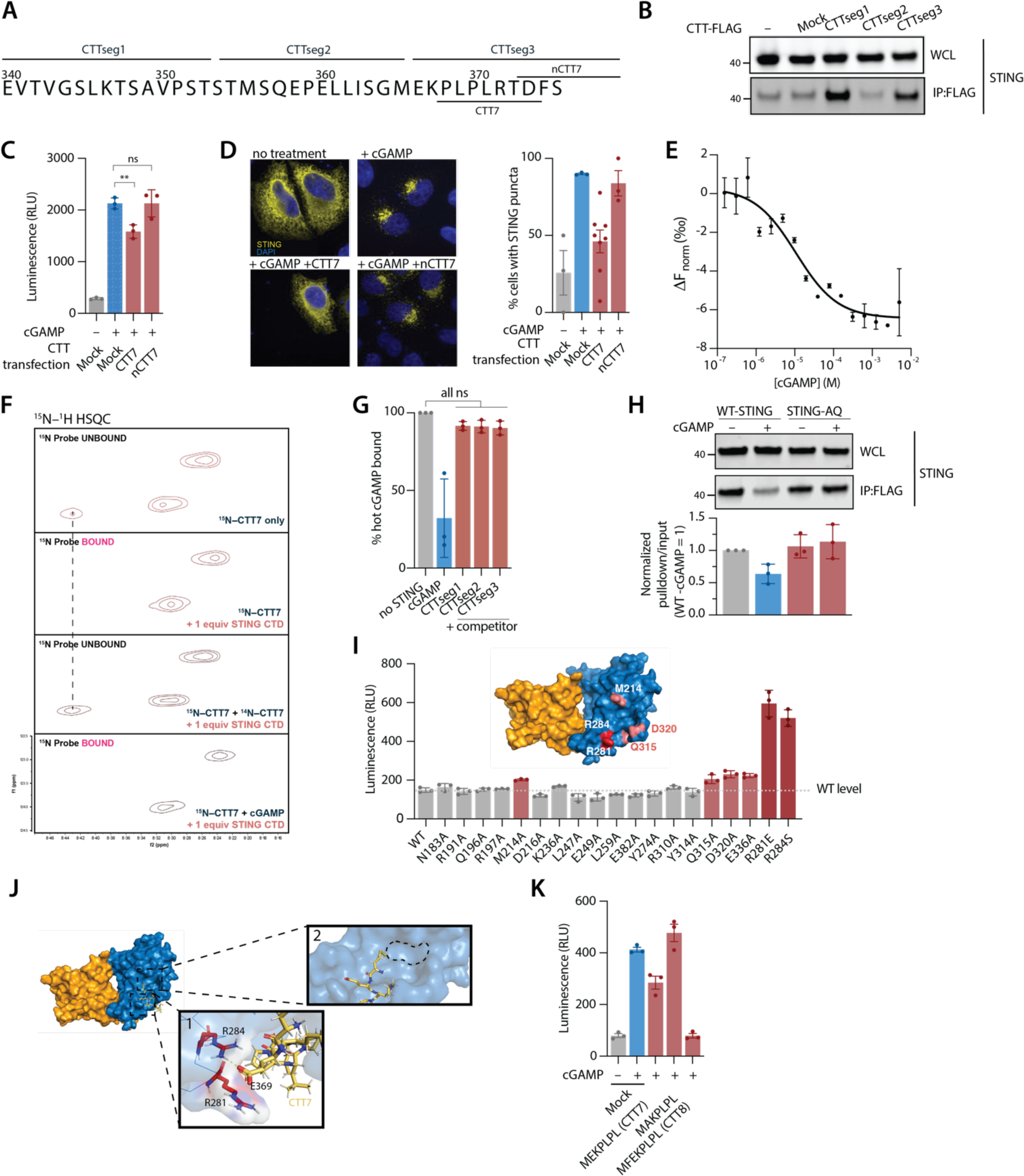
C-terminal tail-derived fragment binds STING to inhibit oligomerization and activation. **(A)** Sequence of STING’s C-terminal tail. Three truncated units (CTT1–3, labelled) were first evaluated for STING affinity. The binding and inhibitory fragment CTT3 was then further truncated into a minimal peptide capable of inhibiting STING signaling (CTT7) and control peptide (nCTT7). **(B)** Co-immunoprecipitation of STING with each of the three truncated units of the CTT in transfected HEK293T cells. **(C)** Relative IFNβ-luciferase activity from HEK293T-Dual reporter cells co-transfected with WT-STING and CTT7-FLAG constructs. **(D)** Left: Representative confocal microscopy images of Hela Kyoto cGAS KO cells co-transfected with WT-STING and CTT7-FLAG constructs; Right: Quantification of percentage of cells with STING puncta under the indicated conditions. **(E)** Binding curve of CTT7 peptide with STING LBD protein measured by microscale thermophoresis. **(F)** ^15^N–^1^H HSQC spectra obtained for 1.5 mM ^15^N-labelled CTT7 alone (panel 1), with one equivalent of STING CTD (panel 2), with one equivalent of both STING CTD and non-isotopically labelled CTT7 (panel 3), and one equivalent each of STING CTD and cGAMP (panel 4). **(G)** Radioactivity assay of ^32^P-labelled cGAMP bound to purified STING CTD in the presence of non-radioactive cGAMP or various fragments of STING CTT (CTTseg1—3). **(H)** Top: Co-immunoprecipitation assay of CTT7 peptide with either WT-STING or STING-AQ, an oligomerization-defective mutant, from transfected HEK293T cells; Bottom: Corresponding quantification of pulldown normalized to input over three independent repeats. **(I)** Relative IFNβ-luciferase activity from HEK293T-Dual reporter cells transfected with a panel of alanine mutants of residues on the STING surface. Constitutively active mutants were determined by higher activity than basal WT indicated by dashed lines and shown in red. **(J)** Molecular docking of CTT7 onto STING CTD. Inset 1 shows a candidate binding site for CTT7 on STING, with potential interacting residues shown. Inset 2 reveals an unoccupied hydrophobic pocket-like site. **(K)** Relative IFNβ-luciferase activity from HEK293T-Dual reporter cells co-transfected with WT-STING with CTT variants. MAKPLPL is a mutant lacking a glutamate residue that potentially interacts with the STING CTD. CTT8 is a lengthened CTT fragment that more potently suppresses STING signaling.

We were unable to obtain co-crystals of STING with CTT7 for structural characterization. Instead, we visualized the interaction of ^15^N-labeled CTT7 peptide with unlabeled STING cytosolic domain (STING residues 138–366) using ^15^N–^1^H HSQC NMR spectroscopy. We were able to detect signals from three of the four NMR-active amidic ^15^N in CTT7 (**Figure 5F**). Although we could not confidently assign the peaks to specific nitrogen atoms, we observed that addition of one molar equivalent of the purified STING cytosolic domain caused a clear upfield shift in one of the ^15^N peaks, suggesting changes to the local environment of the ^15^N-labeled peptide due to binding interactions. This shift was diminished when using STING cytosolic domain containing the R284S mutation that is known to weaken its affinity for the CTT (**Figure S4E**).(*9*) Importantly, addition of the NMR-invisible ^14^N-CTT abolished this shift, indicating direct competition for a shared binding site (**Figure 5F**).

Previous studies hypothesized that cGAMP binding allosterically competes with CTT in binding to STING.(*7*, *9*, *30*) Surprisingly, addition of cGAMP to the ^15^N-CTT7:STING cytosolic domain complex did not change the resultant spectrum (**Figure 5F**). Reciprocally, the addition of CTT7 did not compete with the binding of ^32^P-labelled cGAMP to STING, even at high concentrations of any fragment of CTT (**Figure 5G**), demonstrating for the first time that cGAMP and CTT can bind STING simultaneously. In contrast, using co-immunoprecipitation, cGAMP addition reduced the interaction of CTT7 with WT, but not the oligomerization-defective STING mutant STING-AQ (**Figure 5H**). Together, our data suggest that CTT7 binds STING at the oligomerization PPI interface. It appears that this binding site does not undergo significant conformational changes upon cGAMP binding, since CTT7 can bind equally well to STING with or without cGAMP present, suggesting that this binding site may be easily accessed by antagonists.

To structurally define the CTT7 binding site as a potential druggable pocket, we screened for mutants that no longer bind their own CTT and are therefore constitutively active in the absence of cGAMP. We identified four hyperactive mutants, with three of them (M214A, Q315A and D320A) occurring near residues R284 and R281 implicated in the autoimmune disease SAVI (**Figure 5I**). The fourth residue, E336, is found on the opposite side of the STING body and participates in phase separation-mediated restraint of STING signaling.(*31*) By mapping the first group of residues onto a crystal structure of the apo-STING cytosolic domain (PDB: 4F5W), we identified a candidate binding site for the CTT7 peptide. We computationally docked CTT7 into this site and observed that a salt bridge could form between STING R284 and CTT7 E369 (numbered by its original position in the full-length STING sequence) (**Figure 5J**, inset 2). Indeed, an E369A-CTT7 (MAKPLPL) mutant peptide lacking the negative charge was no longer able to inhibit STING signaling using our interferon reporter assay (**Figure 5K**). Our docking model also predicted that the N-terminal methionine of CTT7 only partially occupies a hydrophobic pocket-like site proximal to the critical R—E salt bridge (**Figure 5J**, inset 1). To further verify this model, we lengthened the CTT with the addition of a phenylalanine residue at position 2 to more fully occupy this hydrophobic pocket. Indeed, the resultant 8-mer (MFEKPLPL) showed a ∼2.7-fold enhancement of affinity for human STING *in vitro* (**Figure S4F**) and fully inhibited STING signaling in cells (**Figure 5K**).

Altogether, our findings demonstrate that context- and cell-type-dependent cysteine post-translational modifications converge on the governance of STING oligomerization: the key mechanistic step that is necessary and sufficient for downstream interferon signaling. In light of this, we determined and took advantage of STING’s natural autoinhibitory mechanism to define a potentially druggable pocket at the oligomerization interface that should provide context-independent control of STING signaling outcomes with future therapeutic development.

## Discussion

In this work, we provide biochemical and functional insights into the roles of cysteine post-translational modifications in signaling activation and inhibition, the mixed STING oligomerization, and its autoinhibition by the CTT, three key aspects of STING biology that are not forthcoming from structural biology efforts to date. We uncovered several new principles that challenge current assumptions and hold major implications that will inform the direction of subsequent development of human STING inhibitors.

We identified a new human STING palmitoylation site at C64 and showed that it plays an essential role in STING activity by preventing basal inactive STING oligomer formation in the absence of stimuli. C64 palmitoylation ensures a transmembrane domain configuration that allosterically controls LBD conformation and ultimately productive TBK1 phosphorylation of IRF3. Interestingly, C64 is the most conserved cysteine within the transmembrane domain of STING across metazoans, including in the sea anemone *Nematostella vectensis* (**Figure S3A**). Sea anemone STING when activated induces autophagy, the primordial antiviral pathway but not the interferon pathway.(*27*) Thus, we are compelled by the possibility that palmitoylation at this site, through organizing the transmembrane domain and STING oligomer assembly, could be important even in the earliest evolutions of metazoan STING signaling. In mammals, STING induces new antiviral mechanisms such as the TBK1–IRF3–interferon axis, gaining modules on the CTT that allow docking of downstream effectors(*32*) and concomitantly conservation of additional cysteines 88, 91 and 148 (**Figure S3A**). In these systems, we showed that C64 maintains importance in regulating STING by allosterically controlling C148, which in turn through formation of disulfides regulates the stability and flexibility of the STING oligomer now required for the productive docking of TBK1 and IRF3.

Our discovery of C64 and C148 as lynchpin sites that regulate human STING signaling prompted us to ask if either residue can be targeted in pursuit of an inhibitor. Unfortunately, our proteomic data indicated that C64 does not possess any reactivity even to the non-specific IA-DTB probe (data not shown), suggesting it is buried and only accessible to enzyme-mediated palmitoylation, an axis of regulation that is more challenging to drug STING-specifically. C148 has recently been targeted for STING inhibitor development. (*17*, *33*) Although there appears to be room for structure-activity relationship analysis for C148-targeting inhibitors, unexpected residue requirements of refined inhibitors suggest that drugging either C64 and C148 is hampered by practical issues of accessibility and specificity.(*33*)

Instead, because we found that cysteine post-translational modifications ultimately converge on controlling STING oligomerization initiation and composition, we sought a strategy that directly modulates the STING–STING interaction necessary for oligomerization. Oligomerization of STING and docking of TBK1 and IRF3 are mutually autoinhibited through sequestering CTT against the STING LBD. In support of this, we showed that an interaction exists between R284 on STING and E369 on the CTT; both residues are conserved in animals with STING capable of inducing the interferon pathway (**Figure S3A**). By examining the mechanism of autoinhibition and CTT release, we identified the minimum 7-mer peptide motif required for CTT-mediated autoinhibition in trans and its binding site on the STING LBD. Although the affinity of CTT7 for STING is modest, its binding site represents a previously unconsidered pocket for inhibition that should, in theory, be amenable to small molecule drug discovery. Already, CTT8 with one extra amino acid residue completely blocks STING signaling. We propose that subsequent efforts to inhibit human STING should directly target oligomerization by screening for binders of the CTT8 pocket, with appropriate counter-screens that exclude binders of the cGAMP pocket and SAVI mutants. Given the increasingly broad implications of unwanted STING activation in autoimmunity and neuroinflammation,(*34*, *35*) it is likely that a STING inhibition campaign, depending on the disease and relevant target cell types—and therefore the relevant configuration of active STING—could necessitate the use of multiple modes of inhibition for most efficacious blockade of aberrant signaling.

## Acknowledgements

We thank April Pawluk and the Arc Institute Scientific Publications Team for constructive feedback on the manuscript. We thank the Peter Kim lab for providing access to peptide synthesis equipment. We thank Martha Cyert and James Ferrell for their helpful discussion and suggestions on palmitoylation research and threshold setting, and all other Li Lab members for their constructive comments and discussion through the course of this study. R.C. thanks the Stanford Graduate Student Fellowship. L.L acknowledges her Stanford startup fund and Arc Institute funding.

## Author Contributions

R.C. and X.C. conceived and coordinated the project, performed experiments, analyzed results, and wrote the manuscript with input from all authors. S.L.E conceived the project, performed experiments and analyzed results. E.N. performed the chemoproteomic experiments and analyzed results. S.R.L. performed the NMR experiment and analyzed results. C.R. analyzed results of RNA-Seq experiments. B.F.C. supervised E.N., provided scientific expertise, coordinated the project, and approved the final manuscript. L.L. conceived the project, coordinated the project, wrote the manuscript, and provided the project funding.

---

**Supplementary Figure 1.**
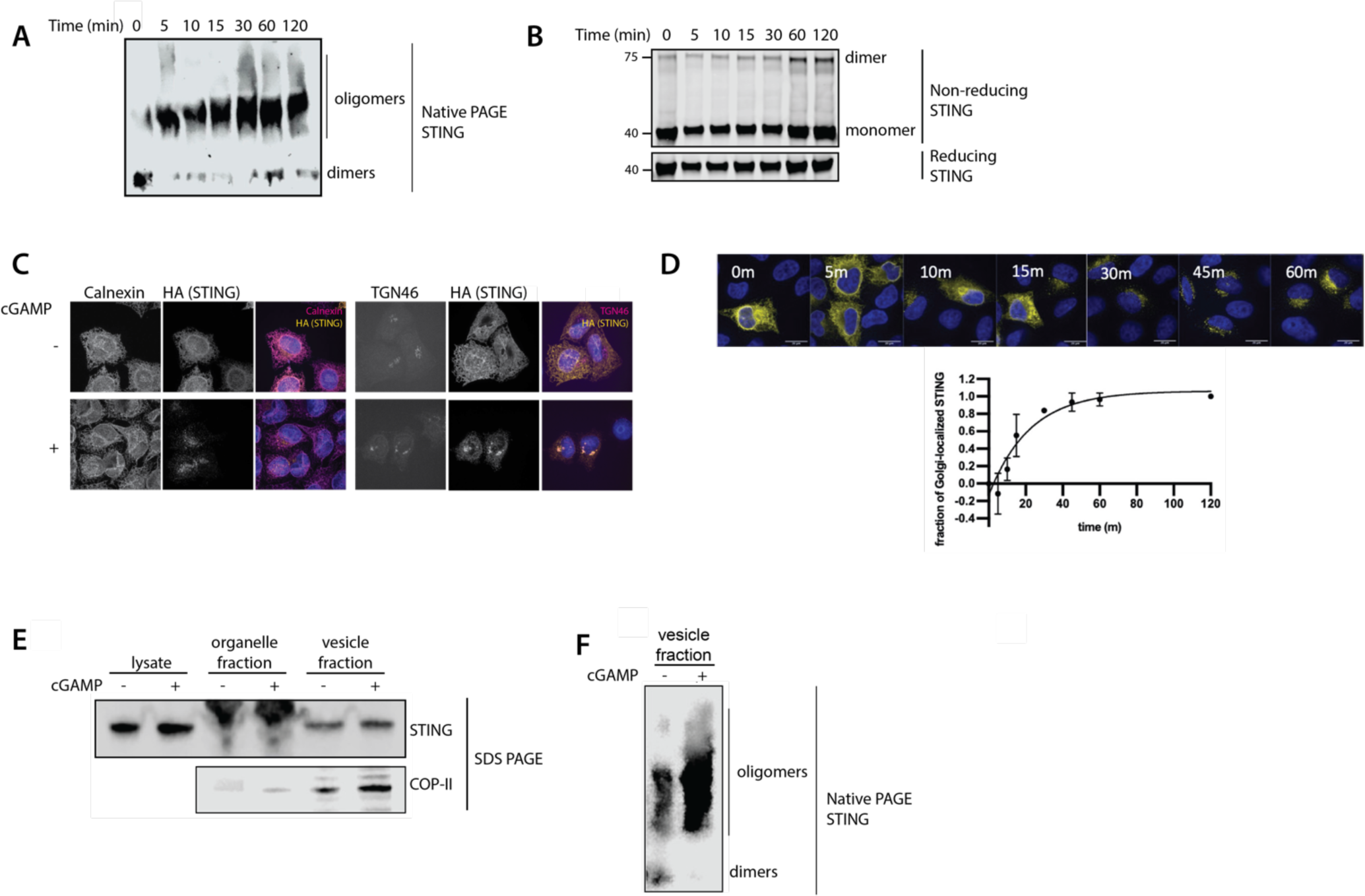
– Related to Figure 1. **(A)** Timecourse of STING oligomerization by Blue-Native PAGE in Hela Kyoto cells expressing STING-HA. **(B)** Timecourse of STING dimer formation as observed by non-reducing SDS-PAGE of HEK293T cells transfected with WT-STING and stimulated with cGAMP for indicated times. **(C)** Confocal microscopy of Hela Kyoto cells expressing WT-STING-HA, showing localization with and without cGAMP. **(D)** (Top) Timecourse of STING trafficking by confocal microscopy in Hela Kyoto cells expressing STING-HA. (Bottom) Quantification showing fraction of Golgi-localized STING at various timepoints. **(E)** SDS-PAGE gel of vesicles enriched through cellular fractionation, as demonstrated by enhancement of COP-II proteins. **(F)** Blue-Native PAGE of vesicle-enriched fraction from (E).

**Supplementary Figure 2.**
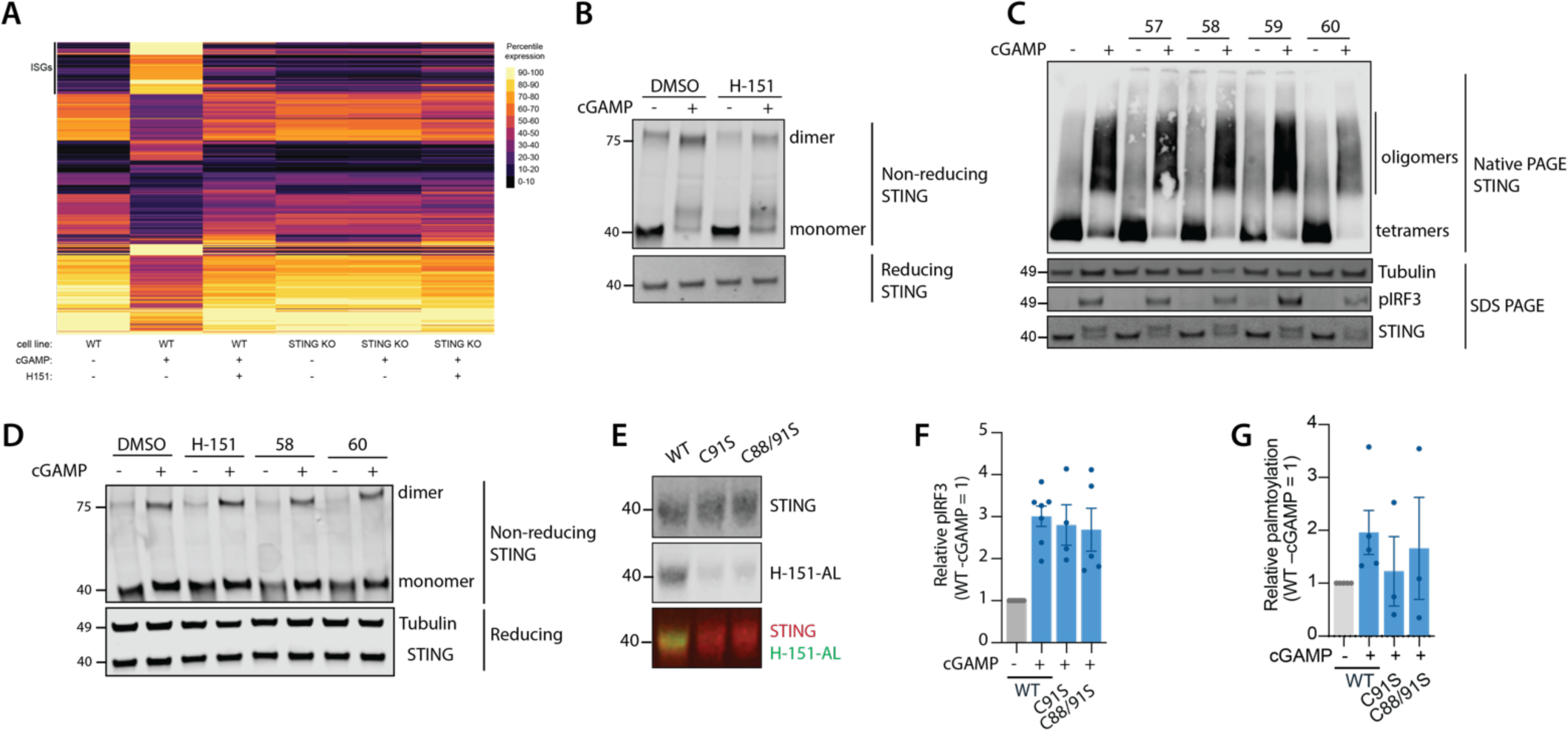
– Related to Figure 2. **(A)** Bulk RNA sequencing of U937 cells treated with 100 μM cGAMP, or both 100 μM cGAMP and 5 μM H-151. **(B)** Representative Non-reducing SDS-PAGE of STING from U937 cells pre-treated with either DMSO or 5μM H-151 for 2 h, then treatment of 100 μM cGAMP stimulation for a further 2 h before lysis. **(C)** Blue-Native PAGE of endogenous STING in Hela Kyoto cGAS KO cells pre-treated with 10 μM of indicated cysteine-reactive compound, followed by 100 μM cGAMP stimulation. **(D)** Non-reducing SDS-PAGE of U937 cells pre-treated with 10 µM of indicated C91-reactive compound followed by 50 μM cGAMP stimulation. **(E)** Alkyne-bearing H-151 (H151-AL) incorporation assay in HEK293T cells expressing STING variants. Cells were labelled in the same manner as palmitoylation assay except H-151-AL was used in place of 17-ODYA. **(F)** Relative pIRF3 levels of HEK293Ts transfected with indicated STING variants followed by cGAMP stimulation. **(G)** Quantification of palmitoylation assays from Figure 2L.

**Supplementary Figure 3.**
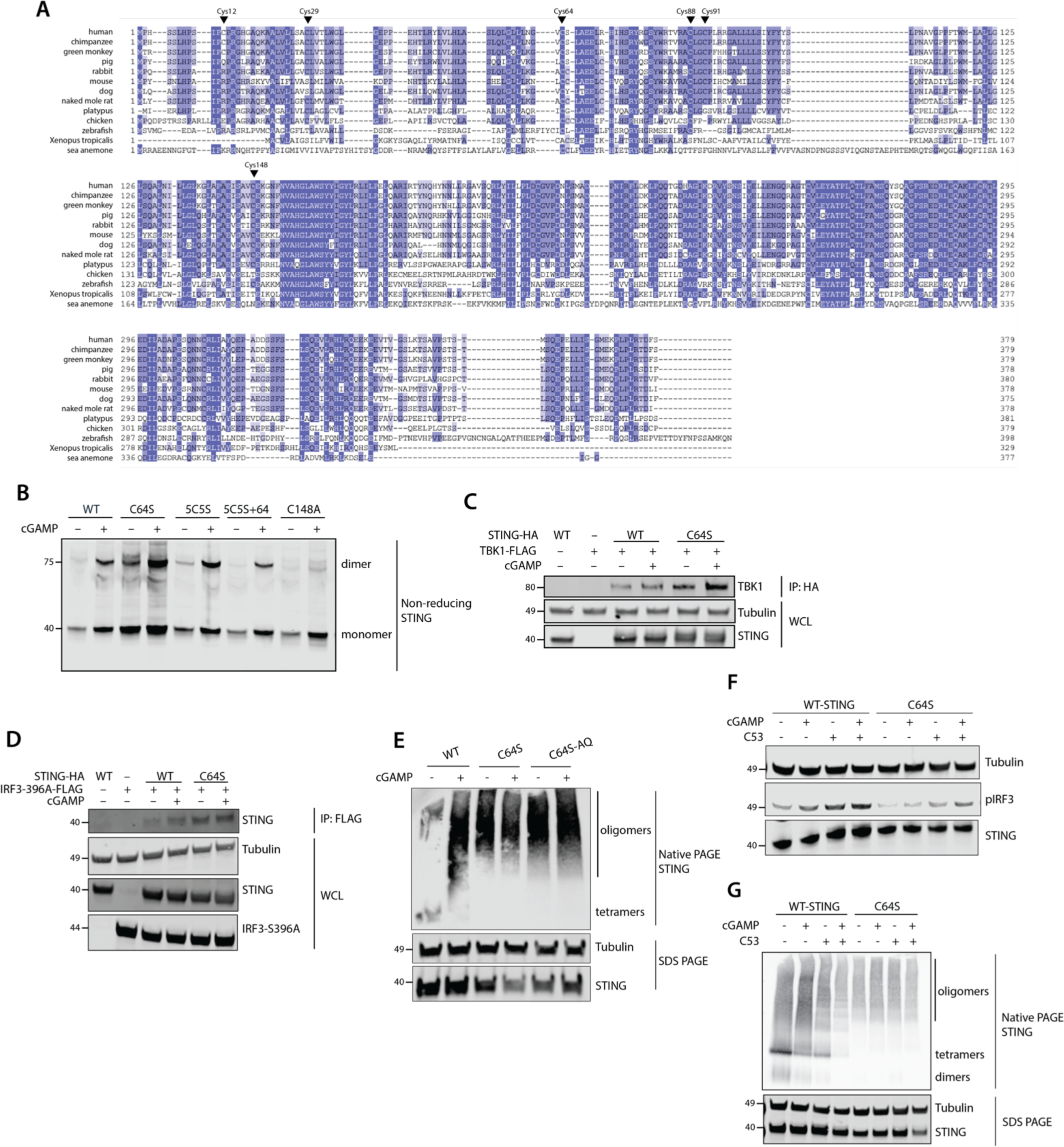
– Related to Figure 3. **(A)** Alignment of the amino acid sequence of STING from various representative species. Sequences of STING were obtained from UniProt, then aligned using COBALT and visualized on JalView. **(B)** Non-reducing SDS-PAGE of HEK293T cells transfected with indicated STING variants followed by cGAMP stimulation. **(C)** Co-immunoprecipitation of TBK1-FLAG with HA-tagged WT-STING or STING-C64S in HEK293T cells. **(D)** Co-immunoprecipitation of IRF3-FLAG (S396A mutant) with HA-tagged WT-STING or STING-C64S in HEK293T cells. IRF3-S386A mutant was used to improve pulldown efficiency. **(E)** Blue-Native PAGE of HEK293Ts transfected with indicated STING variants, including STING-C64S/Q273A/A277Q (C64S-AQ) which combines C64S mutant with an oligomerization deficient mutation pair. **(F)** pIRF3 levels of HEK293T cells transfected with indicated STING variants and stimulated with either 50 μM cGAMP, 10 μM compound C53 or both. **(G)** Blue-Native PAGE of HEK293T cells transfected with indicated STING variants and stimulated with either 50 μM cGAMP, 10 μM C53 or both.

**Supplementary Figure 4.**
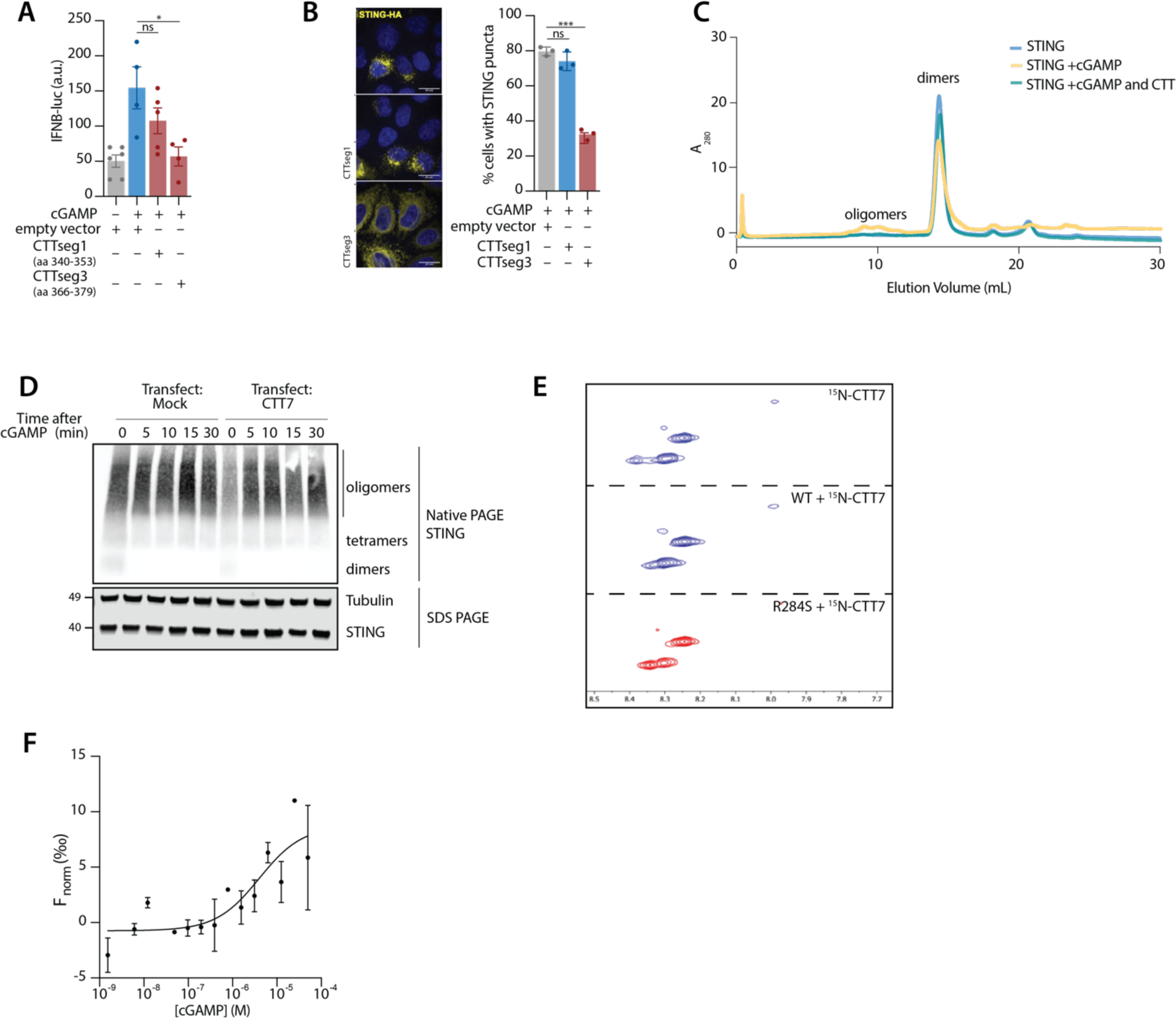
– Related to Figure 5. **(A)** Relative IFNβ-luciferase activity from HEK293T-Dual reporter cells co-transfected with WT-STING and either CTTseg1 or CTTseg3. **(B)** Left: Confocal microscopy of Hela Kyoto cGAS KO cells co-expressing WT-STING-HA with either CTTseg1 or CTTseg3. Right: Quantification showing percentage of cells with STING puncta. **(C)** Size-exclusion chromatography (SEC) of STING LBD with and without addition of 1 μM cGAMP and/or 10 μM CTT peptide. STING LBD was diluted to 1 μM in TBS (pH 8) and incubated with appropriate ligands overnight at 4.C. SEC was performed using TBS (pH 8) as the elution buffer and monitored for protein elution at 280 nm. **(D)** Blue-Native PAGE showing timecourse of WT-STING oligomerization when co-expressed with CTT7 or equal amounts of empty vector DNA in HEK293T cells, after 50 µM cGAMP stimulation for the indicated times. **(E)** Comparison between the ^15^N–^1^H HSQC spectra obtained for 1.5 mM ^15^N-CTT7 alone (panel 1), with one equivalent of STING CTD (panel 2), and one equivalent of STING-R284S CTD. **(F)** Binding curve of CTT8 peptide with STING LBD protein measured by microscale thermophoresis.

## Materials and Methods

| Material | Description | Source and/or Reference | ID |
| --- | --- | --- | --- |
| **Recombinant DNA** |  |  |  |
| pcDNA3-hSTING-HA |  | (Ergun et al., 2019) | N/A |
| pcDNA-hSTING-C148A |  | (Ergun et al., 2019) | N/A |
| pcDNA3-hSTINGΔCTT-HA | | (Ergun et al., 2019) | N/A |
| pcDNA3-hSTINGΔCTT-Q273A/A277Q-HA | | This paper | N/A |
| pcDNA3-CTT1-FLAG (hSTING340–353) |  | This paper | N/A |
| pcDNA3-CTT2-FLAG (hSTING354–366) |  | This paper | N/A |
| pcDNA3-CTT3-FLAG (hSTING367–379) |  | This paper | N/A |
| pcDNA3-CTT3-FLAG-E369R (hSTING367–379) |  | This paper | N/A |
| pcDNA3-CTT7 (hSTING 368–374) |  | This paper | N/A |
| pcDNA3-nCTT7 (hSTING 373–379) |  | This paper | N/A |
| pcDNA3-CTT8 (MFEKPLPL) |  | This paper | N/A |
| pcDNA3-CTT7 E369A (MAKPLPL) |  | This paper | N/A |
| pTB146 hSTING-CTD (aa 137–366) |  | This paper | N/A |
| pcDNA3-hSTING-C29S-HA |  | This paper | N/A |
| pcDNA3-hSTING-C64S-HA |  | This paper | N/A |
| pcDNA3-hSTING-C91S-HA |  | This paper | N/A |
| pcDNA3-hSTING-C88/91S-HA |  | This paper | N/A |
| pcDNA3-hSTING-C12/29/64/88/91S-HA (5C5S) |  | This paper | N/A |
| pcDNA3-hSTING-C12/29/88/91S-HA (5C5S+64) |  | This paper | N/A |
| pcDNA3-hSTING-C12/29/64/88S-HA (5C5S+91) |  | This paper | N/A |
| pcDNA3-hSTING-C12/29/88S-HA (5C5S+64,91) |  | This paper | N/A |
| pcDNA3-TBK1-FLAG |  | Genscript | Cat#: OHu15095D |
| **Chemicals, reagents and peptides** |  |  |  |
| Phusion High-Fidelity DNA polymerase |  | NEB | Cat #: M0530S |
| dNTPs |  | NEB | Cat #: N0447S |
| Phusion GC Buffer |  | NEB | Cat #: B0519 |
| Gibson Assembly HiFi DNA Master Mix |  | NEB | Cat #: E2621 |
| Pfu Turbo DNA polymerase |  | Agilent | Cat #: 600250 |
| Protein A Magnetic Beads |  | Cell Signaling Technologies | Cat #: 73778S |
| Protein A Agarose Beads |  | ThermoFisher | Cat #: 20333 |
| Prolong Diamond Antifade Mountant |  | ThermoFisher | Cat #: P36961 |
| DAPI |  | ThermoFisher | Cat # D3571 |
| DpnI |  | NEB | Cat #: R0176S |
| HisPur Cobalt Resin |  | ThermoFisher | Cat #: 89966 |
| ZymoPure Plasmid Miniprep Kit |  | Zymo Research | Cat #: D4212 |
| ZymoClean Gel DNA Recovery |  | Zymo Research | Cat #: D4002 |
| Direct-zol RNA extraction kit |  | Zymo Research | Cat #: R2050 |
| Protein Labeling Kit RED-NHS 2nd Generation |  | Nanotemper | Cat #: MO-L011 |
| cOmplete protease inhibitor tablets |  | Roche | Cat #: 11697498001 |
| 2',3'-cGAMP | In-house synthesis | (Ritchie et al., 2019) | N/A |
| H-151 | In-house synthesis | (Haag et al., 2018) | N/A |
| H-151AL | In-house synthesis | (Haag et al., 2018) | N/A |
| Peptide H<sub>2</sub>N-MEKPLPL-CO<sub>2</sub>H |  | Genscript | N/A |
| Peptide CTT1<br>H<sub>2</sub>N-EVTVGSLKTSAVP-CO<sub>2</sub>H |  | Stanford Protein and Nucleic Acid Facility | N/A |
| Peptide CTT2<br>H<sub>2</sub>N-STSTMSQEPELLIS-CO<sub>2</sub>H |  | Stanford Protein and Nucleic Acid Facility | N/A |
| Peptide CTT3<br>H<sub>2</sub>N-GMEKPLPLRTDFS-CO<sub>2</sub>H |  | Stanford Protein and Nucleic Acid Facility | N/A |
| 17-Octadecynoic Acid |  | Cayman Chemical | Item #: 90270 |
| BB-CI-Amidine hydrochloride |  | MedChemExpress | Cat #: HY-111347A |
| C53 STING Agonist-12 |  | Cayman Chemical | Cat #: 37354 |
| IRdye 800CW Azide Infrared Dye |  | LI-Cor Biosciences | P/N: 929-60000 |
| AccuPower 2X Greenstar qPCR Master Mix |  | Bioneer | Cat #: K-6251 |
| L-methionine-N-FMOC (<sup>15</sup>N, 98%) |  | Cambridge Isotope Laboratories, Inc. | Cat #: NLM-4632-PK |
| L-leucine-N-FMOC (<sup>15</sup>N, 98%) |  | Cambridge Isotope Laboratories, Inc. | Cat #: NLM-2397-PK |
| L-lysine-alpha-N-FMOC (<sup>15</sup>N<sub>2</sub>, 96-98%) |  | Cambridge Isotope Laboratories, Inc. | Cat #: NLM-4631-0.1 |
| L-proline-N-FMOC (<sup>15</sup>N, 98%) |  | Cambridge Isotope Laboratories, Inc. | Cat #: NLM-3906-0.25 |
| L-glutamic acid-N-FMOC, gamma-tert-butyl ester (<sup>15</sup>N, 98%) |  | Cambridge Isotope Laboratories, Inc. | Cat #: NLM-8960-0.1 |
| **Antibodies** |  |  |  |
| α-tubulin (DM1A), Mouse mAb |  | Sigma-Aldrich | Cat#: T6199; RRID:AB_477583 |
| STING (D2P2F) Rabbit mAb |  | Cell Signaling Technologies | Cat#: 13647; RRID: AB_2732796 |
| HA-Tag (C29F4) Rabbit mAb |  | Cell Signaling Technologies | Cat#: 3724; RRID: AB_1549585 |
| DYKDDDDK (D6W5B) Rabbit mAb |  | Cell Signaling Technologies | Cat#: 14793 RRID:AB_2572291 |
| Phospho-IRF-3 (Ser396) (D6O1M) Rabbit mAb |  | Cell Signaling Technologies | Cat# 10327; RRID: AB_2773013 |
| IRF-3 (D83B9) Rabbit mAb |  | Cell Signaling Technologies | Cat# 4302; RRID: AB_1904036 |
| Phospho-TBK1/NAK (Ser172) (D52C2) Rabbit mAb |  | Cell Signaling Technologies | Cat# 5483; RRID: AB_10693472 |
| TBK1/NAK (E8I3G) Rabbit mAb |  | Cell Signaling Technologies | Cat#: 38066; RRID:AB_2827657 |
| LC3A/B (D3U4C) Rabbit mAb |  | Cell Signaling Technologies | Cat#: 1274; RRID:AB_2617131 |
| Sheep anti-human TGN46 |  | Bio-Rad | Cat#: AHP500GT |
| DYKDDDDK (9A3) Mouse mAb |  | Cell Signaling Technologies | Cat#: 8146 |
| Donkey anti-Sheep IgG (H+L) Cross-Adsorbed Secondary Antibody, Alexa Fluor 647 |  | Life Technologies Corporation | Cat #: A21448 |
| Anti-mouse IgG (H+L), F(ab')<sub>2</sub> Fragment (Alexa Fluor® 488 Conjugate) |  | Cell Signaling Technologies | Cat #: 4408 |
| Goat Anti-Mouse IgG Antibody, IRDye® 680RD Conjugated |  | Li-COR Biosciences | Cat# 926-68070; RRID: AB_10956588 |
| IRDye 800CW Goat anti-Rabbit IgG (H+L) |  | Li-COR Biosciences | Cat# 925-32211; RRID: AB_2651167 |
| **Critical Commercial Assays** |  |  |  |
| QUANTI-Luc |  | Invivogen | Cat #: rep-qlc2 |
| **Software and algorithms** |  |  |  |
| PyMOL |  | The PyMOL Molecular Graphics System, Version 2.5.2., Schrodinger, LLC | https://pymol.org/2/ |
| Maestro |  | Schrodinger | https://www.schrodinger.com/products/maestro |
| GraphPad Prism 9 |  | GraphPad Software | https://www.graphpad.com/ |
| Image Studio Lite |  | LI-COR | https://www.licor.com/bio/image-studio/ |
| **Other** |  |  |  |
| Amicon Ultra-0.5 centrifugal filter unit |  | Merck | Cat#: UFC500396 |
| Amicon concentrators (3 K) |  | Millipore | Cat#: UFC800308 |
| Amicon concentrators (10 K) |  | Millipore | Cat#: UFC901096 |
| Amicon concentrators (30 K) |  | Millipore | Cat#: UFC903024 |
| HiTrap Q Sepharose FF (5 mL) |  | GE Healthcare | Cat#: 17-5156-01 |
| Superdex 200 increase 10/300 GL |  | GE Healthcare | Cat#: 17517501 |
| **Bacterial strains** |  |  |  |
| *E. coli* Rosetta(DE3) cells |  | Sigma-Aldrich | Cat#: 79054 |
| *E. coli* XL1-Blue cells |  | Agilent | Cat#: 200249 |
| **Mammalian cell lines** |  |  |  |
| HEK293T Dual-Null |  | Invivogen | Cat#: 293d-null |
| HEK293 T |  | ATCC | ATCC CRL-3216; RRID: CVCL_0063 |
| U937 |  | ATCC | ATCC ATCC CRL-1593.2; RRID: CVCL_0007 |
| THP-1 |  | ATCC | ATCC TIB-202; RRID: CVCL_0006 |
| HeLa |  | ATCC | ATCC CCL-2; RRID: CVCL_0030 |
| Jurkat |  | ATCC | ATCC TIB-152; RRID: CVCL_036 |
| **Biological Samples** |  |  |  |
| Buffy coat from donors |  | Stanford Blood Center | N/A |

### Material Availability

Further information and requests for reagents should be directed to and will be fulfilled by the lead contact, Lingyin Li.

### Methods

#### Cloning

Unless otherwise stated in subsequent sections, pcDNA-STING plasmids containing point mutations were made by modifying the previously reported wild-type pcDNA-STING plasmid by site-directed mutagenesis using the QuikChange protocol with the indicated primers (Table S1), followed by Dpn1-digestion. All recombinant plasmids were transformed into XL1-Blue competent cells (Agilent) and sequenced for verification.

#### Experimental Models and Subject Details

Human peripheral blood mononuclear cells (PBMCs) were isolated from whole blood using standard Ficoll procedures and were cultured in RPMI (Corning Cellgro) supplemented with 2% human AB serum (Corning) and 100 U/mL penicillin-streptomycin (ThermoFisher). Buffy coat samples from whole blood were obtained de-identified from the Stanford Blood Center so no sex, gender-identify, or age is associated with the sample.

HEK293T, 293T-Dual Null and HeLa Kyoto cells were cultured in DMEM (Gibco) supplemented with 10% FBS (Atlanta Biologics) (v/v) and 100 U/mL penicillin-streptomycin (Gibco) and maintained in 37°C incubators with 5% CO_2_. U937, THP-1 and Jurkat cells were cultured similarly except in RPMI (Gibco) supplemented with 10% heat-inactivated FBS (Atlanta Biologics) (v/v) and 100 U/mL penicillin-streptomycin (Gibco).

#### Cell stimulation

HEK293T cells were plated in a 12-well plate at 300,000 cells/well. After 24h, they were transfected with indicated plasmids using FuGENE 6 transfection reagent (Promega). After another 24h, cGAMP was added to the media at 100 µM. For experiments containing C91-reactive compounds, the cells were pre-treated with these ligands at indicated concentrations for 2 h prior to cGAMP stimulation. After 2 h cGAMP stimulation, cells were lysed directly on the plate in 2x Laemmli sample buffer (LSB) for reducing blots. STING activation was measured by western blot using appropriate antibodies. U937, Hela Kyoto, THP-1, Jurkats, and PBMCs were all stimulated similarly. For non-reducing blots, native lysis buffer (see Blue-Native PAGE) was used to lyse the cells for 15 min at 4°C, before cell debris was pelleted at 14000g for 5 min. The supernatant was mixed with 5x non-reducing LSB containing 50 mM iodoacetamide in place of β-mercaptoethanol and western blot was performed.

#### Blue-Native PAGE

Cells were lysed in native lysis buffer (10% glycerol, 25 mM NaCl, 20 mM HEPES pH 7.0, 1% DDM) supplemented with protease inhibitor tablet (Roche), and solubilized by rotating 30 min at 4°C. Lysates were spun for 15 min at 14000g and supernatant was collected. Lysate was added to 4x native sample buffer (Invitrogen), run on a NativePAGE gel (Invitrogen), and transferred to a PVDF membrane using either a wet transfer system (BioRad) or semi-dry iBlot transfer system (ThermoFisher). Western blotting protocol was followed for membrane staining.

#### Western Blot

Lysates prepared as described in respective protocols were separated on a SurePage Bis-Tris polyacrylamide gel (Genscript) and transferred to a nitrocellulose membrane using the semi-dry iBlot2 system (Invitrogen). The membrane was blocked for 1 h at room temperature (Intercept blocking buffer, Li-COR Biosciences), and incubated with relevant primary antibodies overnight at 4°C. Following three washes in 1xTBS containing 0.1% Tween-20, secondary antibody (anti-rabbit, anti-mouse, Li-Cor Biosciences) was added for 1 h at room temperature, followed by three additional washes in TBS-T. Blots were imaged in IR using a Li-Cor Odyssey Blot Imager.

#### 17-ODYA labeling and click-immunoprecipitation

Cells were transfected with corresponding plasmids as above. 24h following transfection, cells were changed into low-serum DMEM (2% FBS) containing 60 µM 17-ODYA for 3.5 h total, with the addition of 100 µM 1.5 h into this labeling. After labeling and cGAMP stimulation. The cells were washed in cold PBS, then harvested. Cell pellets were lysed in modified RIPA buffer (150 mM NaCl, 4% SDS, 50 mM triethanolamine (TEA) pH 7.4) with protease inhibitors by incubating on rotation at 4°C for 30 min. Lysate was cleared at 14000g for 15 min. Cleared lysate was then diluted five-fold in TEA lysis buffer (1% Triton X-100, 150 mM NaCl, 50 mM TEA pH 7.4) and incubated with prewashed Protein A agarose beads and anti-HA or anti-FLAG antibody on rotation at 4°C overnight. The following day, the beads were washed 3x in TEA lysis buffer. Proteins bound to beads were conjugated to IRdye800-azide in 50 µL PBS with click chemistry reactants for 2 h at 37°C with constant agitation. Click chemistry reactants were freshly prepared as a 5X master mix that consists of 0.5 mM IRdye800-azide, 15 mM BTTAA, 25 mM sodium ascorbate, and 50 mM CuSO_4_·5H_2_O. Beads were then washed 3x in modified RIPA buffer and proteins were eluted by boiling in 1x LSB (from 5x stock diluted in modified RIPA buffer) and analyzed by western blot.

#### Vesicle isolation

HEK293T cells were plated at 4×10^6^ cells per dish in 10-cm dishes. The following day, the cells were transfected with full-length STING. The following day, cells were treated with 200 µM cGAMP for 10 min and then washed in ice cold PBS. The cells were then incubated in ice cold PBS for 5 min and then washed in hypotonic lysis buffer (25mM Tris-HCl, pH 7.5, 50 mM sucrose, 0.5 mM MgCl_2_, 0.2 mM EGTA) and then incubated for 5 min in the same buffer. The plates were then stood vertically for 1 min to remove excess buffer. The cells were then lysed in 2 mL hypotonic lysis buffer with protease inhibitor added. The cells were scraped from the plate and lysed with 30 passes through a 20G syringe. Immediately following lysis the hypotonic buffer was neutralized through addition of hypertonic buffer (2.5 M sucrose, 25 mM Tris-HCl pH 7.5, 0.5 mM MgCl_2_, 0.2 mM EGTA). The lysate was spun at 1000g for 10 min. The supernatant was collected and re-spun at 78000g. The supernatant was again collected and spun at 238000g to collect the vesicles. The pellet was collected, lysed in 2% DDM and run on native page to measure STING oligomerization state.

#### IFN-β Reporter Assay

HEK293T cells stably expressing a secreted IFNβ-luciferase reporter (Invivogen) were transfected with indicated plasmids. After 24 h, cells were stimulated with 100 µM cGAMP. After an additional 20 h, luciferase activity was measured using the Quanti-Luc system (Invivogen).

#### Co-immunoprecipitation

Co-immunoprecipitation experiments were performed as previously described. Briefly, HEK293T were plated and transfected as described above (see Cell Stimulation). For CTT pulldown experiments, transfection was performed with 100 ng pcDNA3-hSTING or a variant and 600 ng pcDNA3-FLAG expressing a CTT fragment. After 20 h, the cells were lysed in 300 µL hypotonic lysis buffer (see Vesicle isolation) supplemented with protease inhibitor tablet (Roche). The lysates were split into two and half was treated with cGAMP. Primary anti-rabbit FLAG antibody (Cell Signaling Technologies) was added (1:50) overnight. Lysates were incubated with Protein A beads pre-washed in lysis buffer for 30 min at room temperature and washed 5 times in hypotonic lysis buffer supplemented with 0.1% DDM buffer. Protein was eluted from beads in 1xLSB and boiled 10 min at 95°C. Protein was analyzed with western blotting. STING–TBK1 pulldown co-immunoprecipitation was performed as previously described (Zhang et al., 2019).

#### Confocal microscopy

HeLa Kyoto cells were plated on glass coverslips and after 24 hours, then transfected and stimulated as described above. Cells were washed once in cold PBS, then fixed for 20 min in 4% paraformaldehyde in PBS. They were then permeabilized for 10 min in 0.1% Triton-X in PBS (PBS-T), and blocked in 3% BSA in PBS-T for 30 min. Primary (Anti-HA, rabbit) was added 1:800 in blocking buffer and incubated overnight at 4°C. Coverslips were then washed three times in PBS-T and incubated with secondary antibodies (A647 anti-sheep 1:2000, A488 anti-mouse 1:1000) for 30 min at room temperature. Following two washes with 1xPBS-T and one wash with 1xPBS, coverslips were mounted on slides using Prolong Diamond Antifade Mountant (ThermoFisher) and allowed to cure for 24 hours. Slides were imaged on both a Leica DM16000 inverted spinning disk confocal microscope as well as a Zeiss LSM Airyscan 980 confocal microscope.

#### Cysteine reactivity profiling

Profile of STING cysteine reactivity was performed using established workflows (Cravatt et al., 2023; Vinogradova et al., 2020). THP-1 cells were treated with either DMSO, 20 µM of compounds **57** to **60**, or co-treated with 100 µM cGAMP for 3 h. Whole native lysates were then chased with iodoacetamide desthiobiotin (IA-DTB). IA-DTB labeled peptides were enriched with streptavidin beads after trypsin digestion and analyzed by MS3 following TMT16plex tagging.

#### Quantitative PCR

Human PBMCs were treated with cGAMP for 16 h. Cells were then lysed and total RNA was extracted using Direct-zol RNA extraction kit (Zymo) following manufacturer protocols. RNA was reverse transcribed in 20 mL reactions containing 500 ng total RNA, 100 pmol Random Hexamer Primers (Thermo Scientific), 0.5 mM dNTPs (NEB), 20 U RNaseOUT (Invitrogen), 1x Maxima RT Buffer (Thermo Scientific), and 200 U Maxima Reverse Transcriptase (Thermo Scientific). Reverse transcription reactions were incubated first for 10 min at 37°C, then for 30 min at 50°C. Reactions were then terminated by incubating for 5 min at 85°C. To quantify transcript levels, 10 mL reactions were set up containing 1x GreenStar Master Mix (Bioneer), 10x ROX dye (Bioneer), 100 nM forward and reverse primers, and 0.7 mL of reverse transcription reactions. To determine Ct values, reactions were run on a Roche Lightcycler 480 II. using the following program: ramp up to 50°C (1.6°C/s) and incubate for 2 min, ramp up to 95°C (1.6°C/s) and incubate for 10 min; then 40 cycles of ramp up to 95°C (1.6°C/s) and incubate for 15 s, ramp down to 60°C (1.6°C/s) and incubate for 1 min. Primer sequences are provided in Table S1.

#### Protein expression and purification

His-tagged STING CTDs were purified as previously described (Ergun et al., 2019). Briefly, Rosetta competent cells transformed with pTB146 vector expressing hSTING CTD lacking CTT3 (hSTING aa 137–366) were grown in 2XYT media with 100 ng/mL ampicillin. Cells were induced at OD 1 with 0.5 mM IPTG and grown overnight at 16°C. Cells were pelleted, resuspended in TBS pH 8, and snap-frozen in liquid nitrogen. Cells were lysed with two freeze thaw cycles, followed by sonication (4 x 30 s at 35% power). Cell debris was removed by ultracentrifugation for 45 minutes at 40,000g. The supernatant was incubated with HisPur cobalt resin (ThermoFisher) for 1 h at 4°C. The beads were washed in 20 mM imidazole in TBS and eluted in 300 mM imidazole. The hisSUMO tag was cleaved overnight at 4°C by Ulp1 (1:1000 molar ratio), and the tag and His-Ulp1 were removed using cobalt resin. Protein was further purified using size-exclusion chromatography on a Superdex 200 Increase 10/300 GL column (GE Healthcare) using an Akta pure system (GE Healthcare) with TBS pH 8 as the running buffer. Fractions containing protein were verified by SDS-PAGE with Coomassie staining and pooled. The protein was concentrated to 10 mg/mL using Amicon 10K concentrators, aliquoted, snap-frozen in liquid nitrogen and stored at −80°C prior to use.

#### cGAMP Binding Assay

cGAMP binding was verified using the radioactivity-based nitrocellulose filter binding assay as previously described (Li et al., 2014). ^32^P-labeled cGAMP was mixed with 100 µM STING CTD and stoichiometric amounts of small molecule competitors (cold cGAMP, CTT1, CTT2 or CTT3, all 10 mM solutions in water). The mixture was loaded into an Amicon Ultra-0.5 centrifugal filter unit and centrifuged at 12,000g for 10 min. For each sample, 5 µL of flowthrough was mixed with 10 mL of scintillation fluid (Research Products International) and readout was performed on a Beckman Coulter LS6500 Liquid Scintillation Counter.

#### Synthesis of ^15^N-labelled CTT7

^15^N isotopically labelled, Fmoc-protected amino acids were obtained from Cambridge Isotope Laboratories and used without further modification. Peptide synthesis was performed on a CSBio CS336X instrument using standard Fmoc-based solid-phase peptide synthesis. 25 μmol Rink amide AM resin (CSBio) was used to initiate the synthesis, with each coupling step performed with 0.1 mmol of protected amino acid for 3 h at room temperature in the presence of HBTU and N,N-diisopropylethylamine (DIEA). Peptide cleavage and removal of the ester protecting group of glutamic acid was performed by stirring the resin in 95% trifluoroacetic acid (TFA), 2.5% water and 2.5% triisopropylsilane for 4 h at room temperature. The peptide was precipitated in ice-cold diethyl ether and collected by centrifugation at 4000*g* for 15 min. The solid was washed twice with cold diethyl ether, then dried over a stream of dry nitrogen gas and redissolved in HPLC-grade water. Purification was performed via high-performance liquid chromatography (HPLC) on a C18 semi-prep column using a water–acetonitrile gradient in the presence of 0.1% trifluoroacetic acid. Fractions were tested by ESI-MS to verify the correct mass of product, collected, pooled and lyophilized.

#### NMR experiments

^15^N-labelled CTT7 prepared as described above was dissolved in D_2_O and reconstituted to 1.5 mM in 1.1x TBS such that the overall solution contained 10% deuterated solvent for signal locking. In samples containing STING, TBS was replaced with buffer containing STING CTD purified as described above and concentrated to 1.5 mM, with the appropriate addition of unlabelled CTT7 (Genscript) or 2’,3’-cGAMP (synthesized in-house according to Ritchie et al., 2019). NMR data were collected on a Varian Inova 600 MHz NMR Spectrometer with a 5 mm room temperature HCN probe. ^15^N–^1^H HSQC data were collected at 25°C using the gNhsqc pulse sequence with 2 scans for 0.085 seconds over 12000 Hz with 2048 complex points in ^1^H dimension, and 200 complex points over 1000 Hz in the ^15^N dimension.

#### Microscale Thermophoresis (MST)

STING CTD was purified as described above, concentrated to 1 µM and lysine labeling was performed using the Red-NHS 2^nd^ Generation Protein Labeling Kit (Nanotemper) according to manufacturer instructions. Labelled protein was eluted with measurement buffer consisting of 50 mM HEPES (pH 7.5), 150 mM NaCl, 0.01% Tween-20 and diluted to 100 nM. Ligands used for measurement (CTT7, CTT8) were prepared as aqueous solutions. Labelled STING CTD was mixed with equal volume of various ligands to reconstitute final concentrations of 50 nM protein and 152 nM–10 mM ligands (in two-fold dilution steps), then incubated for 5 min at room temperature, before MST measurements were recorded on a Nanotemper Monolith using the picoRED channel.

#### Computational modeling of STING–CTT interaction

A monomer of wild-type apo STING C-terminal domain (PDB: 4F5W) was isolated and minimized in Schrödinger Maestro using standard protein preparation workflow using all default settings at pH 7.4, with missing side chains filled against the corresponding FASTA sequence and using the OPLS3e force field. The molecule CTT7 was constructed from a structure file and optimized for all protonation states and stable conformations at pH 7.4 using the LigPrep workflow. Binding site prediction was run and a site containing the residues R284, R281, D320 and Q315 was selected and used to generate a receptor grid suitable for peptide docking (all other default parameters). The receptor grid was used to dock all prepared conformations of CTT using Glide at standard precision.

---

**Table S1:**
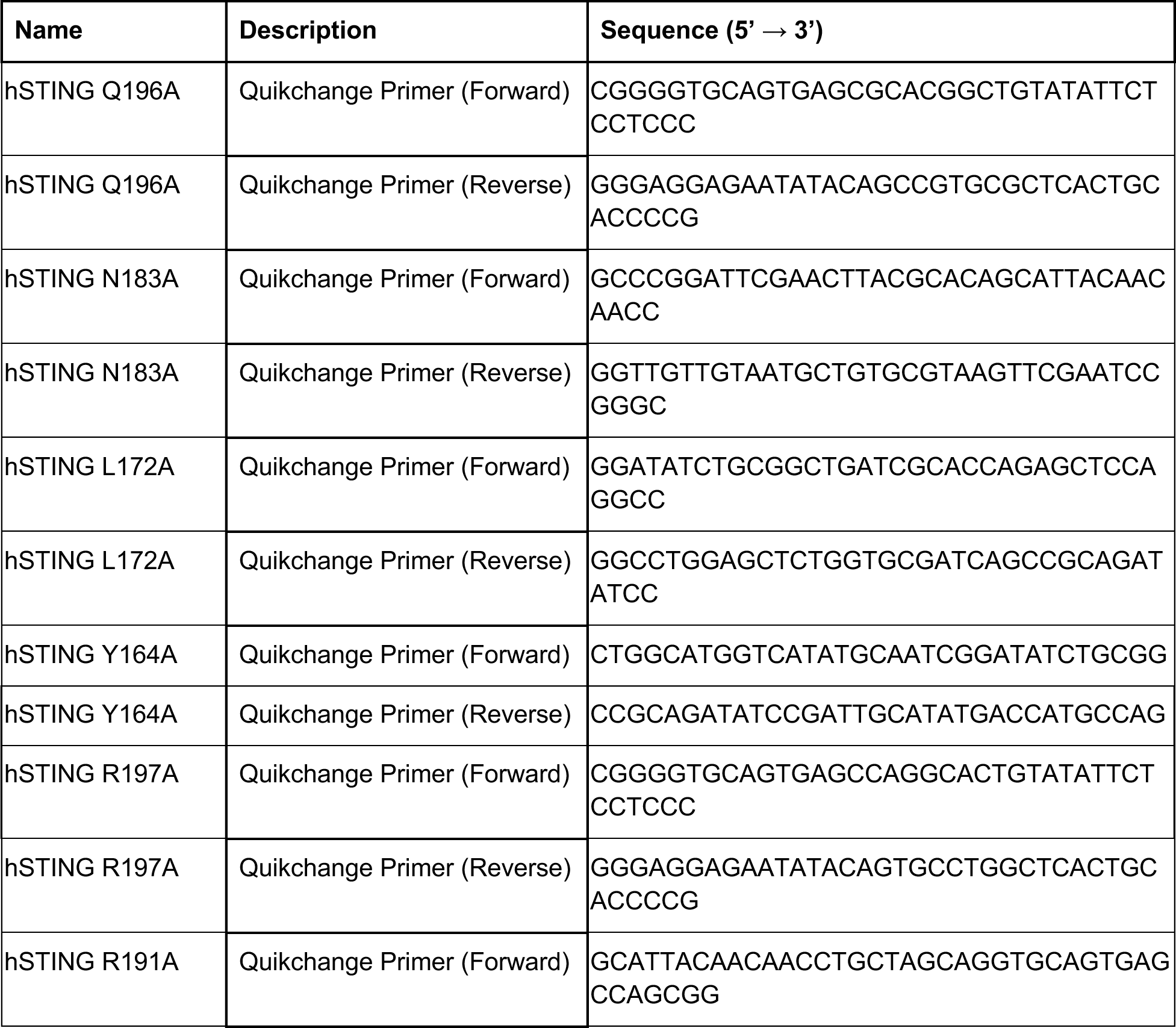
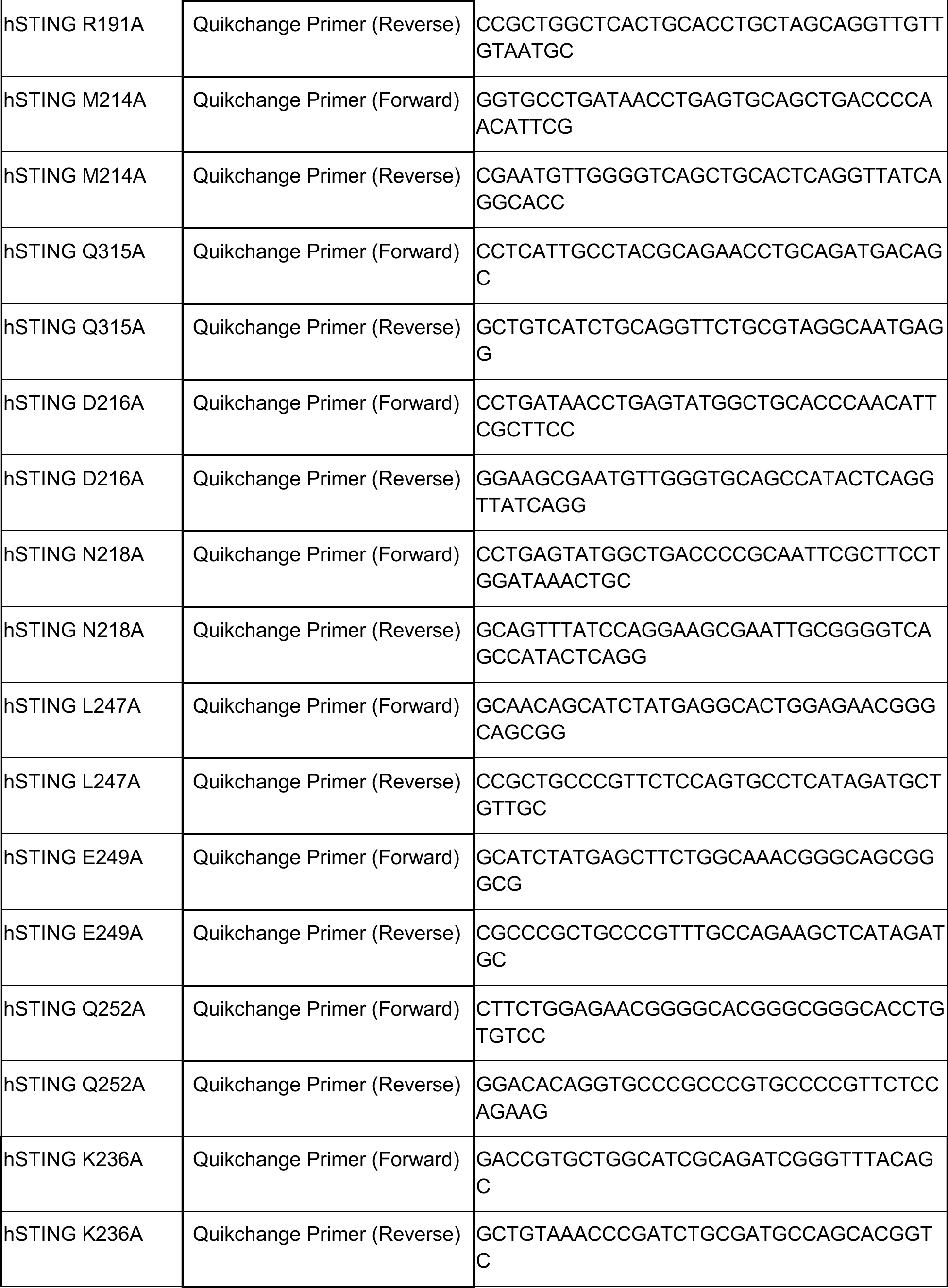
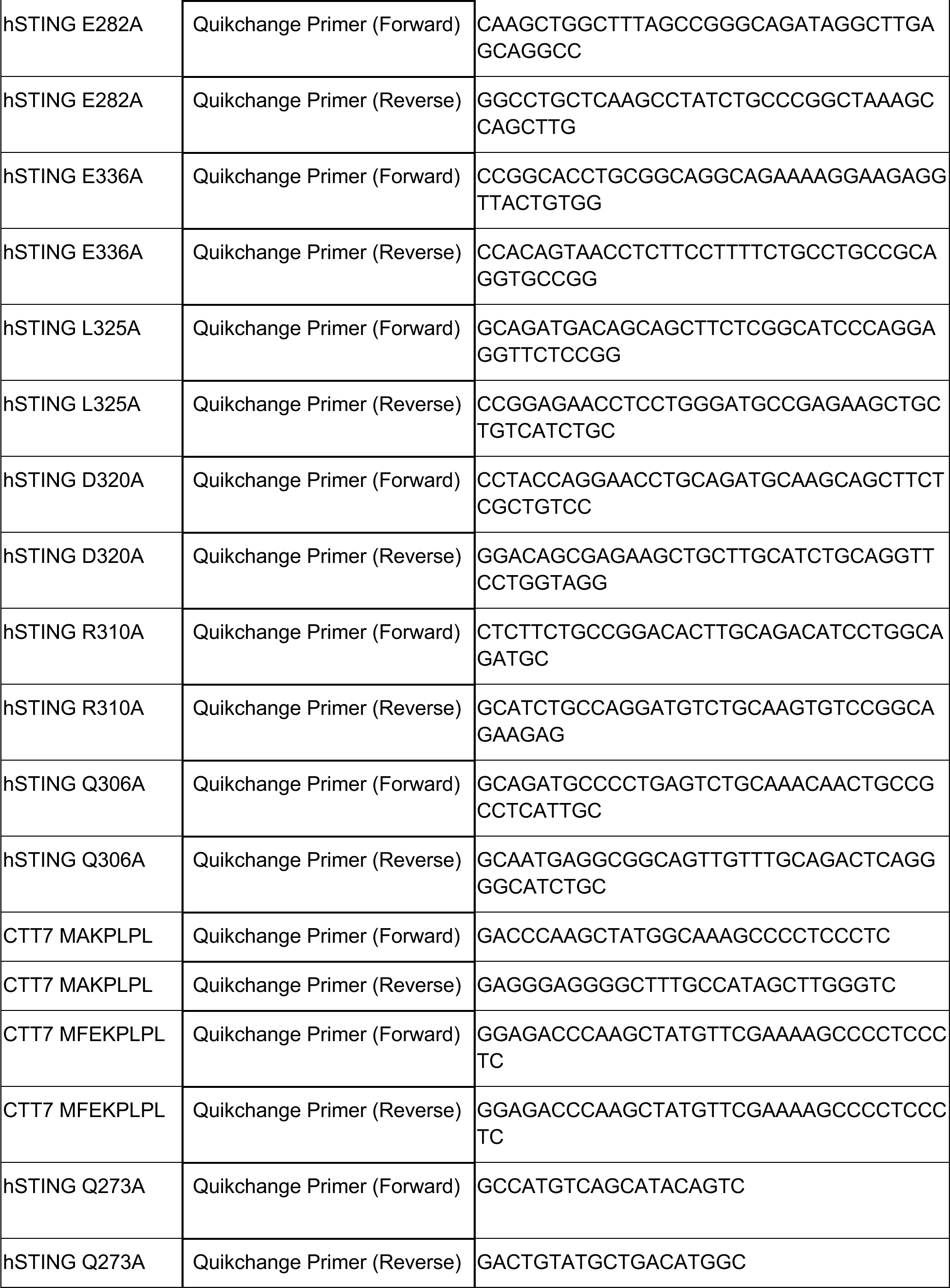
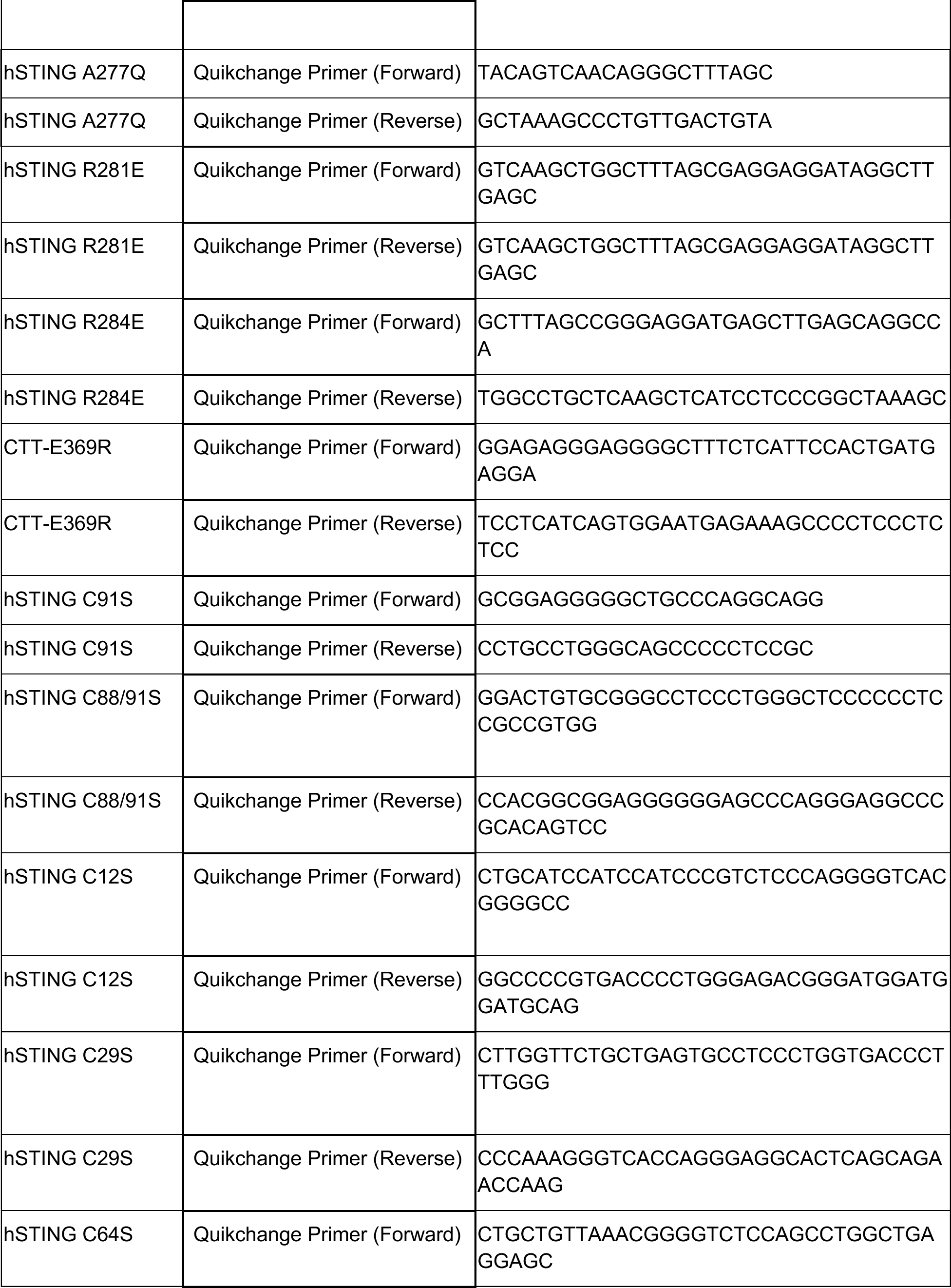
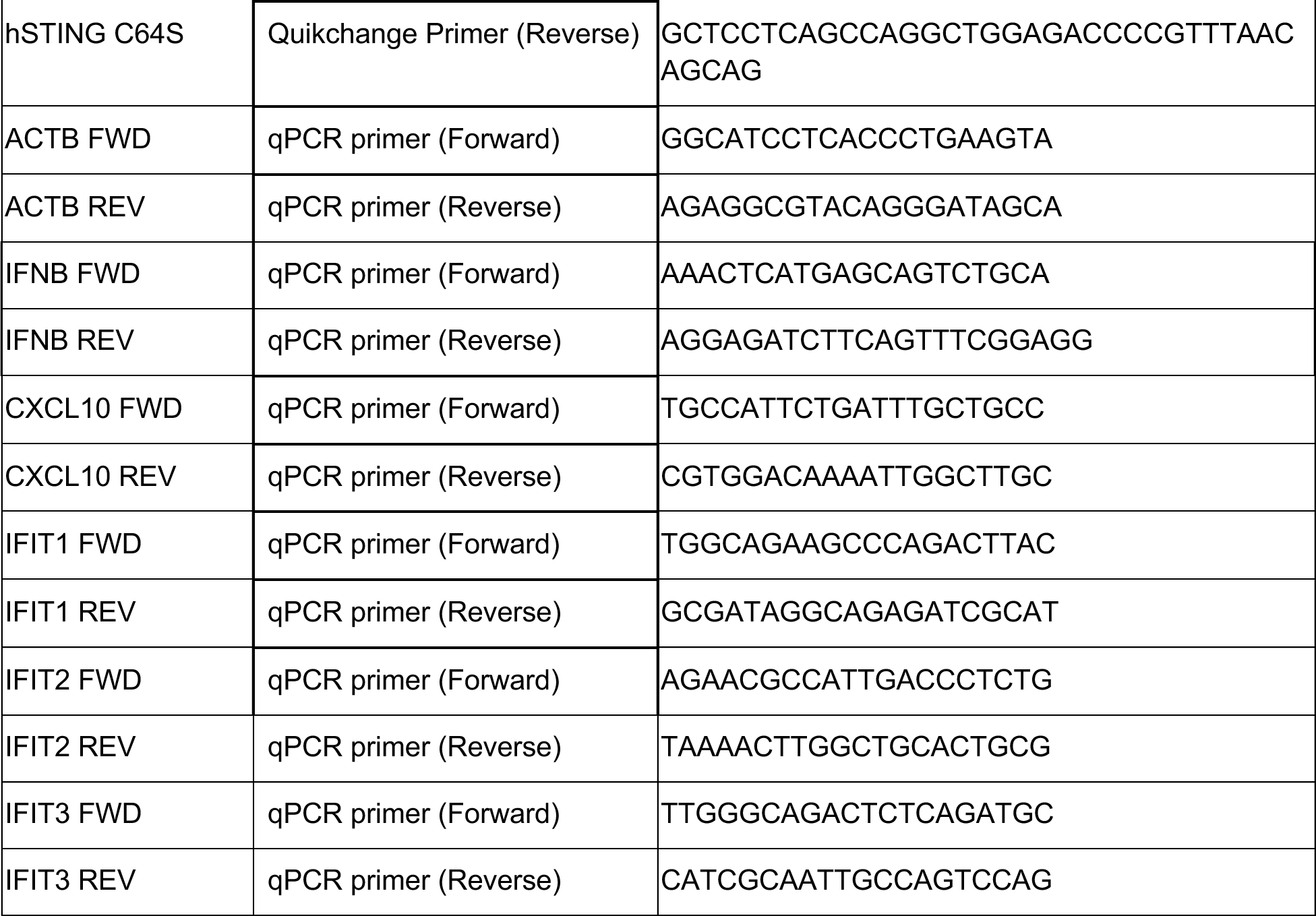
Oligonucleotide sequences used in this study.

